# Microbial Nanocellulose Biotextiles for a Circular Materials Economy

**DOI:** 10.1101/2021.09.22.461422

**Authors:** Theanne N. Schiros, Romare Antrobus, Delfina Farías, Yueh-Ting Chiu, Christian Tay Joseph, Shanece Esdaille, Gwen Karen Sanchiricco, Grace Miquelon, Dong An, Sebastian T. Russell, Adrian M. Chitu, Susanne Goetz, Anne Marika Verploegh Chassé, Colin Nuckolls, Sanat K. Kumar, Helen H. Lu

## Abstract

The synthesis and bottom-up assembly of nanocellulose by microbes offers unique advantages to tune and meet key design criteria—rapid renewability, low toxicity, scalability, performance, and degradability—for multi-functional, circular economy textiles. However, development of green processing methods that meet these criteria remains a major research challenge. Here, we harness microbial biofabrication of nanocellulose and draw inspiration from ancient textile techniques to engineer sustainable biotextiles with a circular life cycle. The unique molecular self-organization of microbial nanocellulose (MC) combined with bio-phosphorylation with a lecithin treatment yields a compostable material with superior mechanical and flame-retardant properties. Specifically, treatment of MC with a lecithin-phosphocholine emulsion makes sites available to modulate cellulose cross-linking through hydroxyl, phosphate and methylene groups, increasing the interaction between cellulose chains. The resultant bioleather exhibits enhanced tensile strength and high ductility. Bio-phosphorylation with lecithin also redirects the combustion pathway from levoglucosan production towards the formation of foaming char as an insulating oxygen barrier, for outstanding flame retardance. Controlled color modulation is demonstrated with natural dyes. Life cycle impact assessment reveals that MC bioleather has up to an order of magnitude lower carbon footprint than conventional textiles, and a thousandfold reduction in the carcinogenic impact of leather production. Eliminating the use of hazardous substances, these high performance materials disrupt linear production models and strategically eliminate its toxicity and negative climate impacts, with widespread application in fashion, interiors and construction. Importantly, the biotextile approach developed in this study demonstrates the potential of biofabrication coupled with green chemistry for a circular materials economy.

---

The linear economy that has been the dominant production model since the Industrial Revolution drives climate instability and threatens ecological and human health. The textile industry, in particular, is reliant on industrial agriculture for cellulosic fibers,^1^ nonrenewable petrochemicals to produce synthetic fibers, dyes, tanning and finishing agents, and chemically and energy intensive processing.^2^ It is responsible for 10% of global carbon emissions,^3^ 20% of global waste water,^4^ 35% of marine microplastic pollution,^5^ and is expected to take up 25% of the global carbon budget by 2050.^6^ For instance, the cattle industry is the single leading driver of deforestation,^7^ while chrome tanning of leather creates large scale mutagenic, teratogenic, and carcinogenic chemical pollution.^8^ Leather alternatives created from petrochemical-based textiles and coatings have emerged. However, these materials are not biodegradable and shed nano- and micro-plastics, which contain endocrine-disrupting chemicals and further absorb and accumulate persistent organic pollutants as they move through ecosystems. Microplastics have been found in the intestines of marine animals and humans, in human blood,and, recently, in human placenta, and have been observed to cross the blood-brain barrier.^9-10^ Thus, there is a pressing need for new fabrication strategies that minimize and close material and energy loops in a regenerative, circular economy, to make industrialization compatible with sustainable development goals.

This challenge has motivated a search for more sustainable, bio-based textiles, especially leather alternatives,^11^ including mycelium and pineapple leather, and gene editing of yeast to produce collagen. Microbial nanocellulose (MC) is a highly crystalline biopolymer produced extracellularly by bacteria, one of the most promising in terms of cellulose yield being *Acetobacter xylinus*.^12^ Under aerobic conditions, these bacteria biosynthesize extracellular cellulose nanofibrils (10–100 nm in diameter) that self-assemble into a three-dimensional layered pellicle with high water content (>98%)^12^ at the air-media interface under static conditions.

Although identical in chemical structure to plant-based cellulose, MC is distinctly characterized by its readily extractable nanofiber network, degradability, excellent tensile strength due to high degrees of polymerization and crystallinity (80-90%), and the possibility to control these and other physical properties including porosity during biosynthesis.^13^ Importantly, MC can offer a rapidly renewable raw material while eliminating the land, water and chemicals usage of production of agricultural^14^ and wood pulp cellulose.^15^ For instance, the amount of cellulose produced by eucalyptus in a 10,000 m^2^ area of land over 7 years could be achieved at higher purity by microbial fermentation in a 500 m^3^ bioreactor in 22 days.^16^

However, like many naturally occurring biopolymers, including the aforementioned bio-based leather alternatives, the hygroscopicity of as-fabricated MC results in unstable mechanics that hinders translation to textile applications.^17^ Strategies to improve the mechanical and moisture stability of “bioleathers” typically involve heavy metals and/or synthetic plasticizers and coatings which compromise material biodegradability, introduce human and ecological toxicity, and increase flammability,^18^ making development of innovative green processing a major research priority.

Yet, for millenia, societies around the world have created durable, water-repellent leather using non-toxic tanning methods, including brain and organ tanning followed by smoke tanning, dating back at least 5,000 years to “Ötzi” (Iceman).^19^ Unfortunately, much of these ancient techniques have been lost, and the chemical mechanism of fat tanning is not fully understood. It has been proposed that braining and smoking hides are forms of aldehyde tanning, in which the aldehyde carbon reacts with the amine nitrogen of collagen to produce an imine product, which reacts with an amine on a neighboring polypeptide to covalently cross-link the collagen into leather.^20^ Brain tissue is high in fatty acids and lecithin. The lipid portion of the lecithin molecule polarizes away from water in the emulsion, while the phosphate and choline groups are attracted to it, stabilizing the tanning emulsion and facilitating interaction between the oils and collagen.

Inspired by these ancient practices, we hypothesize that lecithin phosphatidylcholine will modify the cross-linking and material properties of MC through its hydrophilic [-OH] groups to make it suitable for use as a bioleather. An increase in ductility of MC with application of traditional brain and smoke tanning processes provided a positive control to further encourage this line of investigation (Figure S1). Following a bottom-up approach of microbial biosynthesis of nanocellulose coupled to processing wih a lecithin emulsion, we have created a high performance bioleather that exhibits high tensile strength, ductility, outstanding flame retardance, and soil biodegradability (Figure 1). Here, a symbiotic colony of *Acetobacter* and yeast (SCOBY) metabolizes sugars into cellulose nanofibers via enzymatic pathways.^13^ A variety of carbon sources, such as glucose, fructose, and sucrose, which may be extracted from agro-industrial by-products, can be utilized for scalable production of MC (Movie S1). A range of colors and design motifs are achieved using historically important natural dyes and waste-to-resource strategies, offering a non-toxic alternative to the brown color resulting from the Maillard reaction. Tailoring the shape of the cultivation vessel to the pattern geometry enables zero waste production, as shown for our bioleather bag prototypes in Figure 1. The microbial bioleather created here demonstrates key steps of a circular material economy, including renewability, low toxicity, functional and environmental performance, biodegradability and scalability.

**Figure 1.**
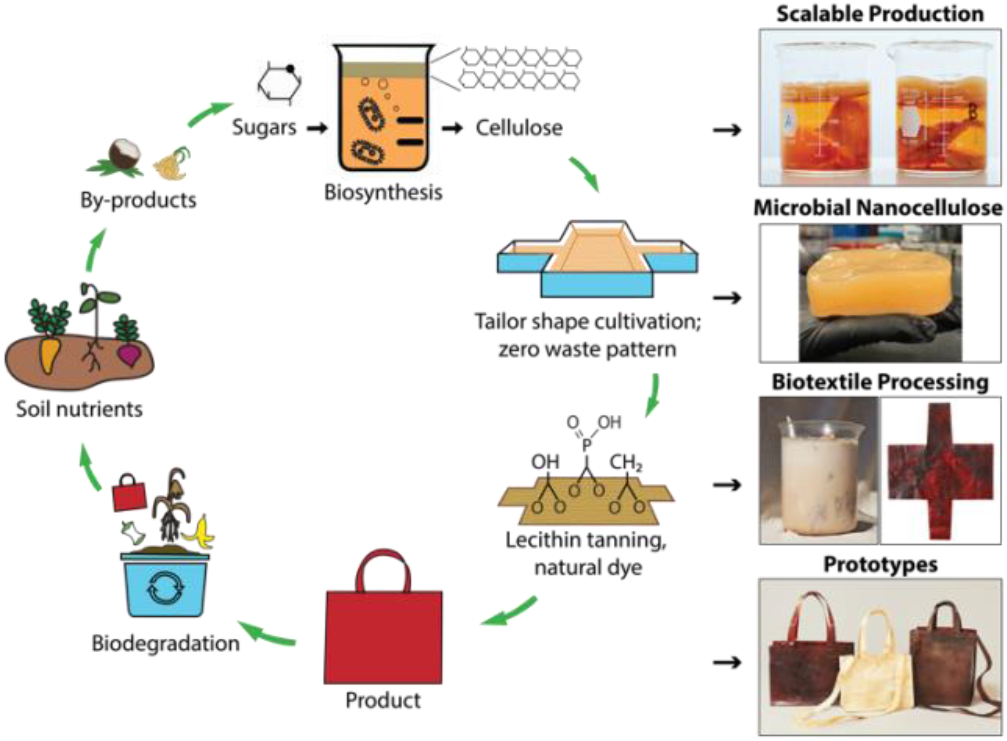
Microbial biofabrication and green processing of minimal waste products with a closed-loop life cycle (left). Right: Scalable MC production from different carbon sources, including fructose (beaker A) and sucrose (beaker B); lecithin tanning and biocoloration--of textiles biosynthesized to the geometry of a design pattern, to create products, such as our bioleather bag prototypes, that meet performance and biodegradability criteria for a circular economy.

We investigate the effect of lecithin treatment on MC and compare with an aldehyde smoke treatment, as documented for brain tanning of hides in order to complete the conversion to leather. As-fabricated, lecithin “tanned”, aldehyde tanned (smoked), and lecithin and aldehyde tanned MC are denoted MC, LT, S, and LTS, respectively. A dense, three-dimensional network of unaligned, intertwined nanofibers is observed with scanning electron microscopy (SEM) on MC surfaces before and after processing (Figure 2A). Treatment with lecithin and aldehyde tanning, separately and sequentially, yields no significant differences in fiber size relative to as-fabricated MC (average diameter of 71.00 ± 19.00 nm, Table 1). These observations indicate that the emulsion does not coat the fibers but is rather removed with rinsing. However, compositional analysis with energy dispersive x-ray measured nearly four times higher phosphorous concentration for LT than the MC control (2.02% vs. 0.57%, ^p<0.05, n=3), supporting the hypothesized favorable interaction between the phosphocholine group of lecithin with cellulose (Figure S2).

**Figure 2.**
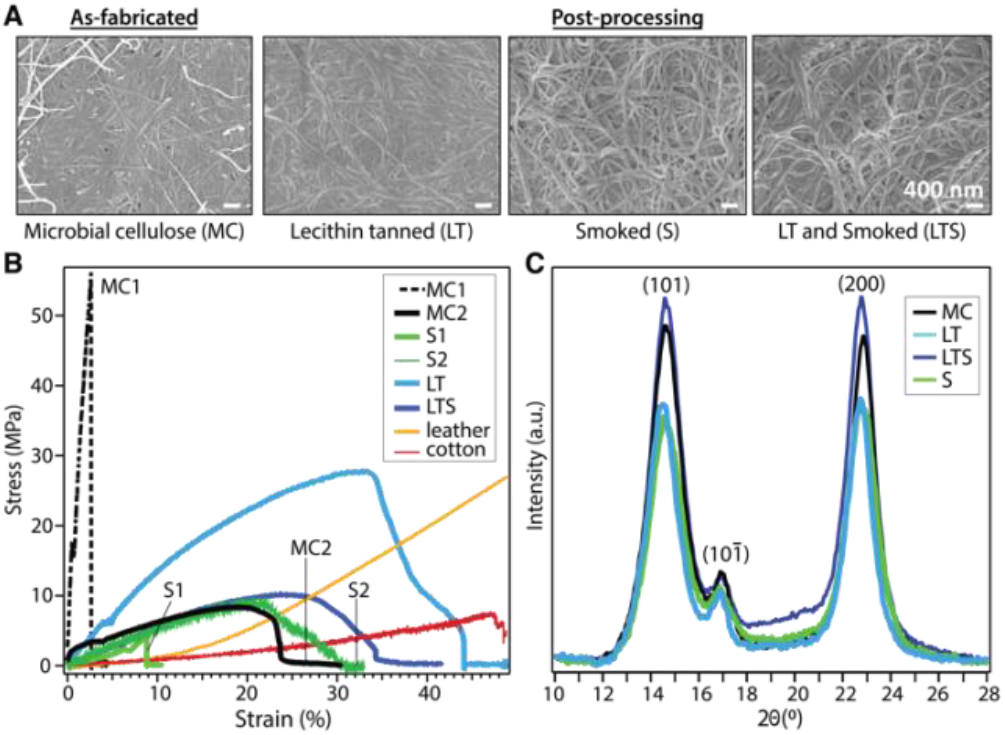
Characterization of MC Biotextile physical properties. (A) nanofiber morphology (SEM); (B) tensile properties; and (C) crystallinity (XRD) of microbial cellulose (MC), MC treated by lecithin (LT) and/or smoke tanning (LTS, S). Sample batches (MC1 and MC2, S1 and S2) were produced from the same biosynthesis and processing conditions; the variation reflects the influence of MC hygroscopicity on mechanical properties which is overcome with lecithin tanning (LT).

**Table 1.**
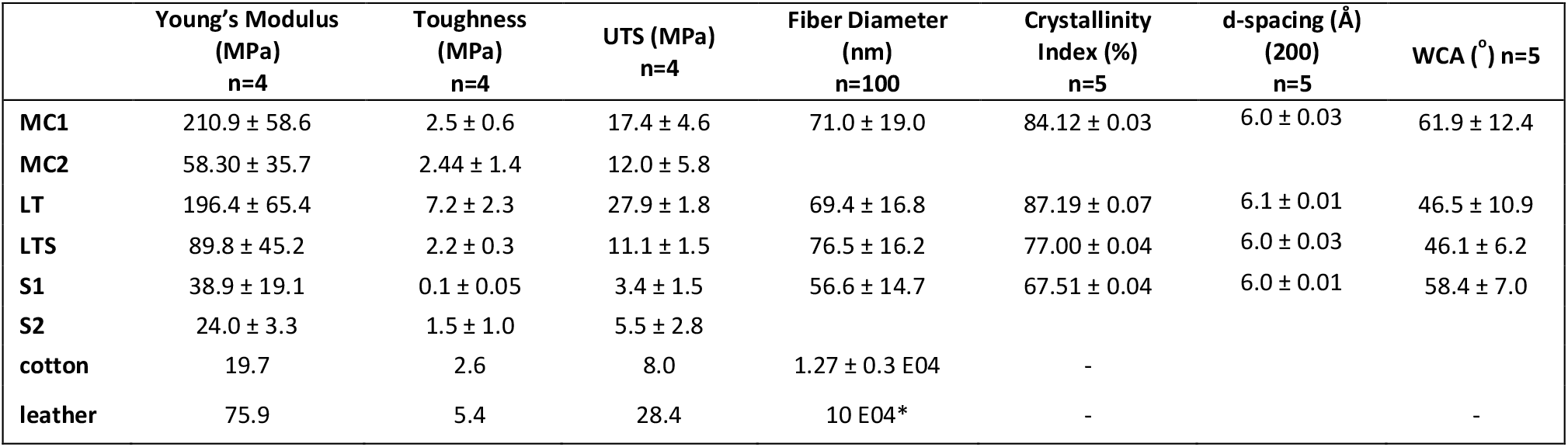
Physical properties of as-fabricated and treated biotextiles compared to cotton and leather. Mechanical properties, including Young’s modulus, toughness and ultimate tensile strength (UTS) (n=4), fiber diameter (n=100), crystallinity index and lattice (d-)spacing (n=5) obtained from x-ray diffraction, and water contact angle (WCA, n=5)) for as-fabricated (MC) and processed (LT, S, LTS) samples shown in Fig. 2. *Fiber diameter for leather based on published data.^41^ ^ indicates significance between groups (*α* = 0.05, n=5). All quantitative values are reported as means ± standard deviation, with n equal to the number of replicates per group, with significance attained at p<0.05.

Common to unprocessed biopolymers, as-fabricated MC samples exhibit diverse stress-strain profiles under uniaxial tensile testing (Figure 2B). The typical trade-off between tensile strength and ductility, the latter a reflection of flexibility which considers both the strain at yield point and toughness, is apparent for dried MC. Despite the same biosynthesis and processing parameters, MC exhibits elastic moduli ranging from 58.30 ± 35.71 to 210.91 ± 58.61 MPa for MC1 (high tensile strength, brittle) and MC 2 (lower tensile strength, greater ductility), respectively (Table 1). Without lecithin treatment, smoked (S) samples also have variable mechanical properties. Lecithin tanning stabilizes the mechanical properties and increases the elastic modulus, toughness, and maximum stress (^ p<0.05, n=4) of the material. The LT group exhibits higher tensile strength than leather or cotton up to strain values of 33% and 45%, respectively (Figure 2B). Surprisingly, the superior mechanical properties of LT are not reflected in the bulk microstructure as probed with x-ray diffraction (Figure 2C). The (1-10), (110) and (200) Bragg peaks of cellulose I are largely unaffected by lecithin tanning, with only a minor increase in crystallinity index (87% vs 84%, Table 1), calculated using a peak deconvolution method,^21^ and unchanged d-spacing and similar crystalline domain size relative to MC (Table S1). A similar increase in tensile strength and ductility unaccompanied by changes in microstructure has been observed for phosphorylated cellulose with low phosphorus concentrations comparable to those measured for LT.^22^

Flame-retardance is an important design criterion for performance textiles, but industrial flame-retardant chemicals are considered hazardous substances linked to autoimmune diseases, neurological and reproductive problems, birth defects, and cancer.^23^ Since phosphorous compounds are effective in reducing biopolymer flammability,^24^ phosphorylation with LT prompted investigation of MC biotextile flame retardance. Remarkably, when exposed to a direct 2054°C flame, LT biotextiles do not ignite (Figure 3A). After 40 seconds, a control aluminum brazing rod is introduced and melts within seconds of application of the flame, while the biotextile continues to deflect the flame, without propagation. When LT samples are repeatedly exposed to the direct flame until surface char is formed, the bulk LT biotextile is observed to be intact just below the easily removable surface char, underscoring its outstanding flame-retardant capacity (Figure 3A).

**Figure 3.**
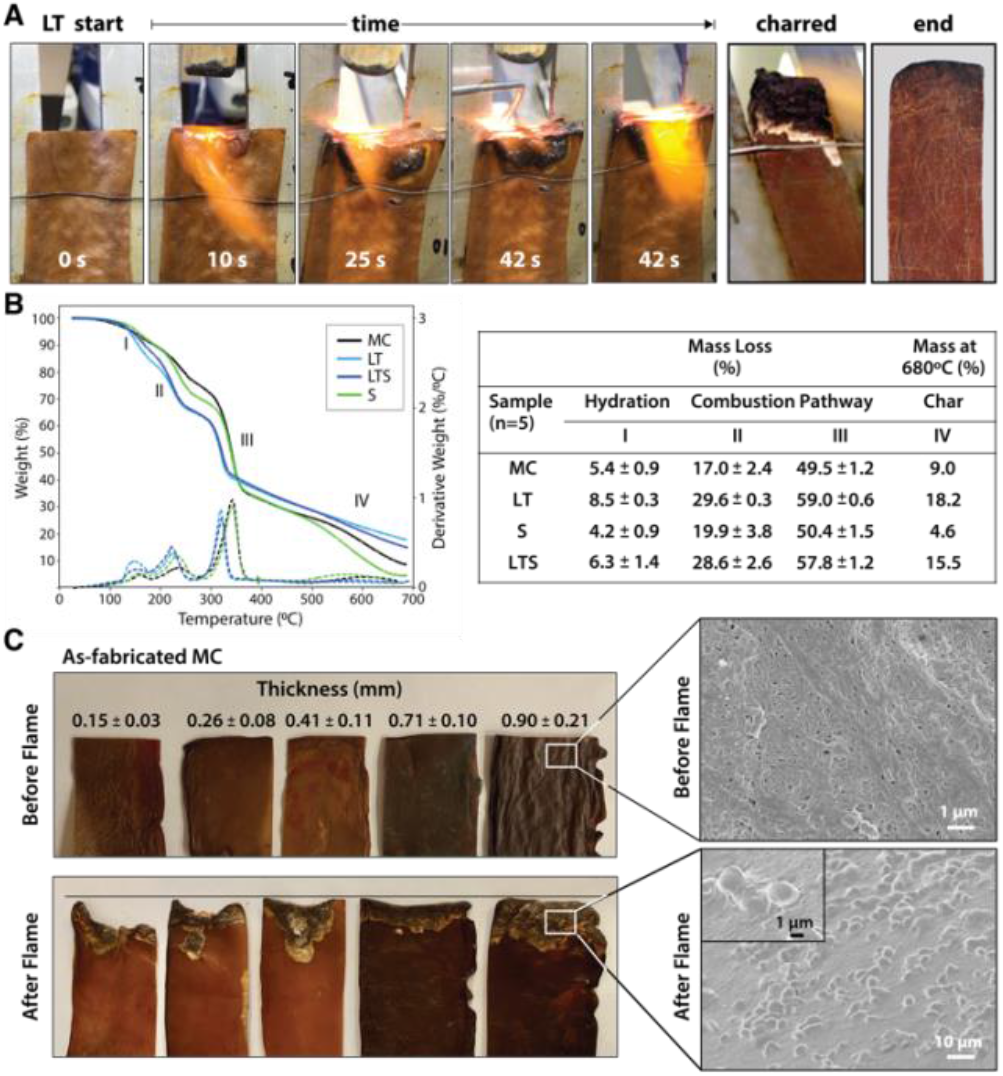
Flame Retardance of MC Biotextiles. (A) Under a 2054°C flame, lecithin-tanned (LT) microbial cellulose does not ignite and deflects the flame from the surface, while an aluminum brazing rod introduced at 39 seconds promptly melts under the flame. After repeated exposure until char formation is observed (charred), LT biotextiles remain intact under the surface ash (end). (B) Thermogravimetric analysis (TGA) curves (left) and data (right) of as-fabricated (MC) and treated (LT, S, LTS) biotextiles under nitrogen atmosphere shows the phospholipid treatment directs the combustion chemistry toward ash formation over the glucosan formation to promote flame retardance. (C) The nanofiber morphology and layered microstructure of as-fabricated MC biosynthesized to sufficient thickness (2.5 cm in hydrated state) also demonstrates flame retardance, evidence by resistance to material lost to combustion. (D) SEM micrograph showing surface morphology before (top) and in charred region after (bottom) flame testing; the torched region of the latter shows a morphology typical of a flame-retardant material.

Analysis of the thermal stability and decomposition behavior of as-fabricated and treated biotextiles provide insights into the mechanism of flame retardance. Thermogravimetric analysis (TGA) and dTGA curves (Figure 3B) show a three-step mass loss, representing: (I) the initial evaporation of free and bound water between 25-200°C; (II-III) polymer decomposition (210-240°C); and (IV) the production of either pyrolysis-based levoglucosan or flame-resistant char (300-360°C). Higher mass loss is observed for LT relative to MC in regions I-III: 8.50±0.25% vs 5.38±0.87% at 150 °C (region I); 29.55±0.25 vs 16.96±2.37% at 232 °C (region II) and 58.99±0.55% vs 49.51±1.21% at 340°C (region III). Finally, a lower decomposition temperature onset is also observed for LT, and aliphatic compounds are decomposed into char with a greater residual mass for LT than MC at 680°C (18.18% vs 8.99%).

The data indicates that lecithin treatment lowers the decomposition temperature of MC and redirects the combustion pathway from formation of levoglucosan towards formation of foaming char as an insulating oxygen barrier, resulting in the outstanding flame resistance. This is attributed to the addition of phosphorous compounds with lecithin tanning, which promotes dehydration to yield unsaturated compounds that polymerize at high temperatures to a cross-linked, intumescent foaming char. Notably, as-fabricated MC also demonstrates a degree of flame retardance, which may be a consequence of the nanoscale self-organization of MC fibers into a layered structure, visible under the deflected flame (Figure 3A). To investigate the influence of the layered microstructure on flame retardance, as-fabricated MC samples of varying thickness (obtained by removing layers from a hydrated MC pellicle (45 × 54 cm, 2.5 cm thick, Figure S3)), dried under ambient conditions, are flame tested (Figure 3C). The mass loss to combustion decreases with increasing MC thickness until 2.5 cm, beyond which point MC material loss is minimal, although greater than for LT biotextiles of the same thickness (Figure 3A, C).

The formation of intumescent bubbles on the nanofiber surface exposed to the flame is observed in the charred portion of LT, as well as for as-fabricated MC of sufficient thickness (≥ 25 mm and 0.9 mm in hydrated (pellicle) and dehydrated states, respectively), in high magnification SEM images (Figure 3D). The multi-layer structure of MC mimics that of a multilayer nanocoating, well known as physical surface barriers that limit heat and oxygen transfer between the fire and cellulose, while the densely packed arrangement of MC nanofibers further limits oxygen available to support a flame. Bubble formation is accompanied by generation of water vapor, which favors production of an insulating foaming char barrier, that in turn, prevents combustion by reducing heat transmission to the MC surface, and inhibiting oxygen and combustible volatiles from diffusing to the flame.

To further investigate the effect of the layered structure of MC on flame response, we attempted to disrupt it by blending MC into a slurry and casting it into a film. We note that the MC nanofiber morphology drives self-assembly into a hydrogen-bonded sheet during dehydration without chemical intervention. Surprisingly, while combusting more readily than as-fabricated MC, cast MC films also demonstrate a degree of flame resistance. Electron micrographs of the cross-sections of the films reveal that blended MC *self-organizes to recover the original layered structure* with the nanofiber morphology intact (Figure S4). Thus, in contrast to the ignitable behavior of plant-based cellulose, the unique nano- and microscale assembly of MC, and in the case of LT, modified chemical properties, imbue the material with superior and non-toxic flame retardance.

Molecular level insights on the chemical origin of the improved mechanical properties for LT biotextiles are revealed by x-ray photoelectron (XPS) and Fourier transform infrared (FTIR) spectroscopy. A peak centered at 134 eV in the phosphorous (P 2p) XPS (Figure4A) indicates that lecithin tanning introduces phosphorus in oxidized states. Carbon 1s XPS reveals an increase in carboxyls, esters and anhydrides, observed at binding energies of ∼286.6 (C-O-H), 288.1 (C-O-C), and 289.1 eV (–COOR), respectively.^25^ These chemical groups, along with P=O (530.9 eV) and C-O-P bonds (533.5 eV), ^26^ also modify the O 1s XPS after lecithin tanning (Figure 4A).

**Figure 4.**
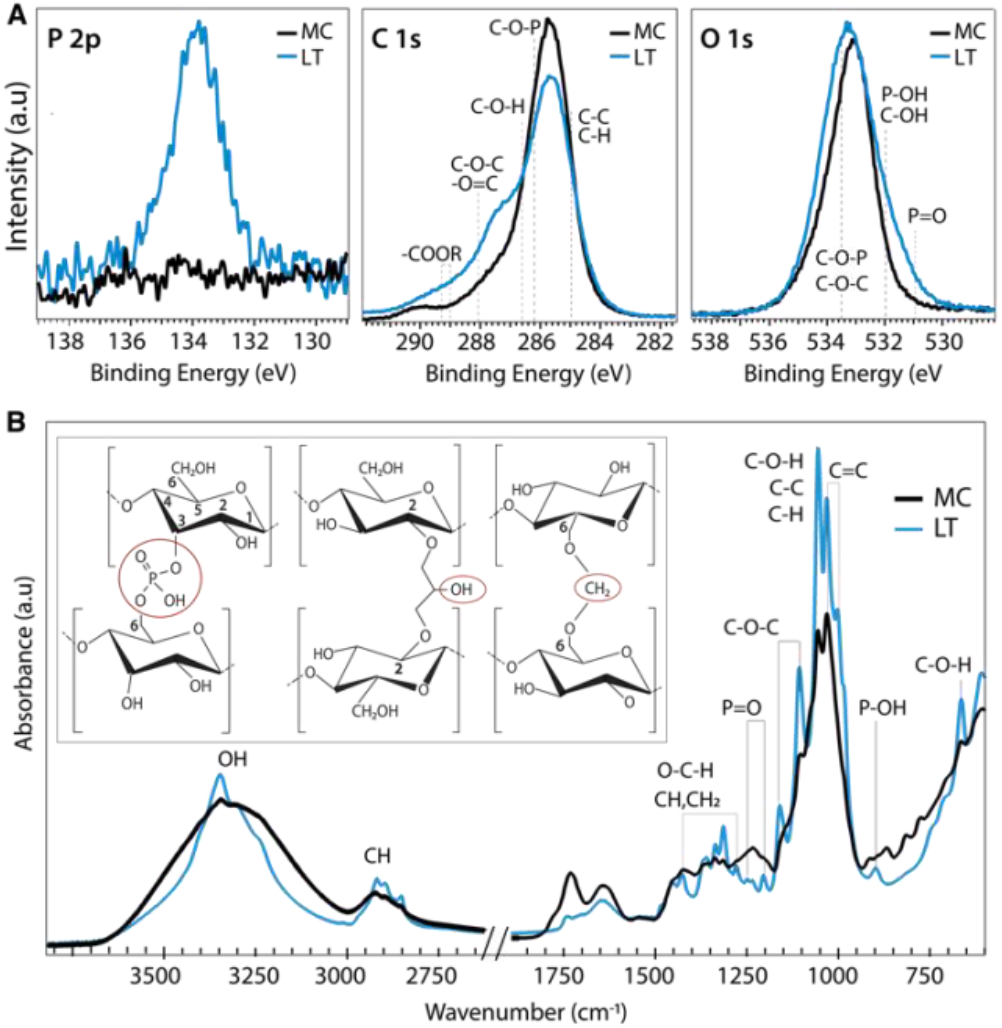
Atomic and Molecular-Level Chemical Analysis. (A) X-ray photoelectron spectroscopy (XPS) and (B) Fourier Transform Infrared (FTIR) spectra of microbial cellulose textiles before (MC) and after lecithin tanning (LT). **Inset**: schematic depiction of proposed cross-linking mechanism with phosphate, methylene and carboxyl group bridges (left to right) for LT.

Chemical fingerprints of bonding observed with FTIR provide mechanistic insights on modulation of cellulose cross-linking with lecithin tanning (Figure 4B). New peaks representative of phosphated cellulose appear for LT, including P=O resonances at 1207.4, 1230 and 1252 cm^1^, and a -P-OH stretch vibration at 898 cm^−1^.^26-28^ Further, enhancement of -OH vibrational peaks characteristic of the cellulose skeleton at 3450-3000 cm^1^ and 664 cm^−1^, and methyl and methylene resonances, including C–H vibrational bands at 1437-1245 cm^−1^ and 3000-2853 cm^−1^ and the C–H deformation (1428, 1370 and 1316 cm^−1^), CH_2_ (1184-1104 cm^−1^) and C-O-H groups (690 cm^−1^) is observed for LT relative to MC.^27-29^ FTIR peak ratios, expressed as the Lateral Order Index (LOI),^30^ shows a higher degree of overall order for LT compared to MC throughout the bulk of the material (Table S4, Figure S5).

Collectively, the data suggests that the lecithin-phosphatidylcholine treatment modifies cellulose cross-linking through phosphate, exocyclic CH_2_ and carboxyl groups at well-defined binding sites; proposed mechanisms for these chemical groups are shown schematically (left to right, Figure 4B inset). Methylene groups may form upon reaction with the hydroxyl group at the 6 position (Figure 4B inset), which can react ten times faster than the other -OH groups.^31^ Further, carboxyl groups observed with XPS can form ester bonds with hydroxyl groups, which are stronger than the hydrogen bonds of the unmodified cellulose fibers. Low phosphorous concentrations (Figure S2) and unchanged lattice parameters (Table S1) indicate far fewer links are made than hydrogen bonds broken.

Indeed, probing the hydrogen-bond intensity in cellulose with FTIR absorbance ratios at different wavelengths, using the C-H stretching band as an internal standard,^32^ shows a clear decrease number of intermolecular hydrogen bonds for LT (Methods, Table S2). This supports our claim that chemical cross-links through methyl, phosphate, and carboxyl bridges form at the expense of hydrogen bonding physical cross-links. This implies that the net strength is higher because of these tighter cross-links (Figure 4B inset), which is confirmed by the higher average modulus (196.4 vs 134.6 MPa) for LT vs MC. The stabilization and improvement of mechanical properties with LT may therefore be attributed to disruption of intrinsic hydrogen bonds between cellulose chains at reactive hydroxyl, methyl and phosphate group sites, leading to an increased interaction between cellulose chains that effectively transfers and distributes applied stress. Additionally, the unprecedented flame retardance promoted by incorporation of phosphates may also be enhanced by the addition of methyl groups with lecithin tanning, which can decrease the rate of combustion by promoting cross-linking during pyrolysis.

Color is a feature vital to the commercial success of textiles, and one which currently comes at a high cost of toxicity through use of synthetic dyes.^2.4^ Natural dyeing is one of the most ancient of arts, dating back to ∼4000 BC Mesopotamia. Leveraging the high biosorption capability of MC,^12^ we color microbial biotextiles with natural dyes, including important historical dyes, such as madder, cochineal, and indigo, as well as those extracted from waste, including discarded onion skins and nails, which provide tannins and iron acetate, respectively, that bind plant chromophores to cellulose and modify color (Figure 5). To eliminate the additional water and energy demands of dip dyeing, the dye bath and culture media can be combined to achieve coloration during biosynthesis (Figure S6). Surface treatment with heated soy wax adds texture and water repellency to the biotextiles (Figure 5.15-16, Movie S2). Surface design motifs, including texture and spatial control in coloration, utilizing ancient folding and printing techniques, denoted Shibori and Hapa Zome in Japan and Adire in West Africa, are demonstrated (Figure 5.17, 5.22), as well as our bio leather sneakers and wallet prototypes (Figure 5.21-5.22).

**Figure 5.**
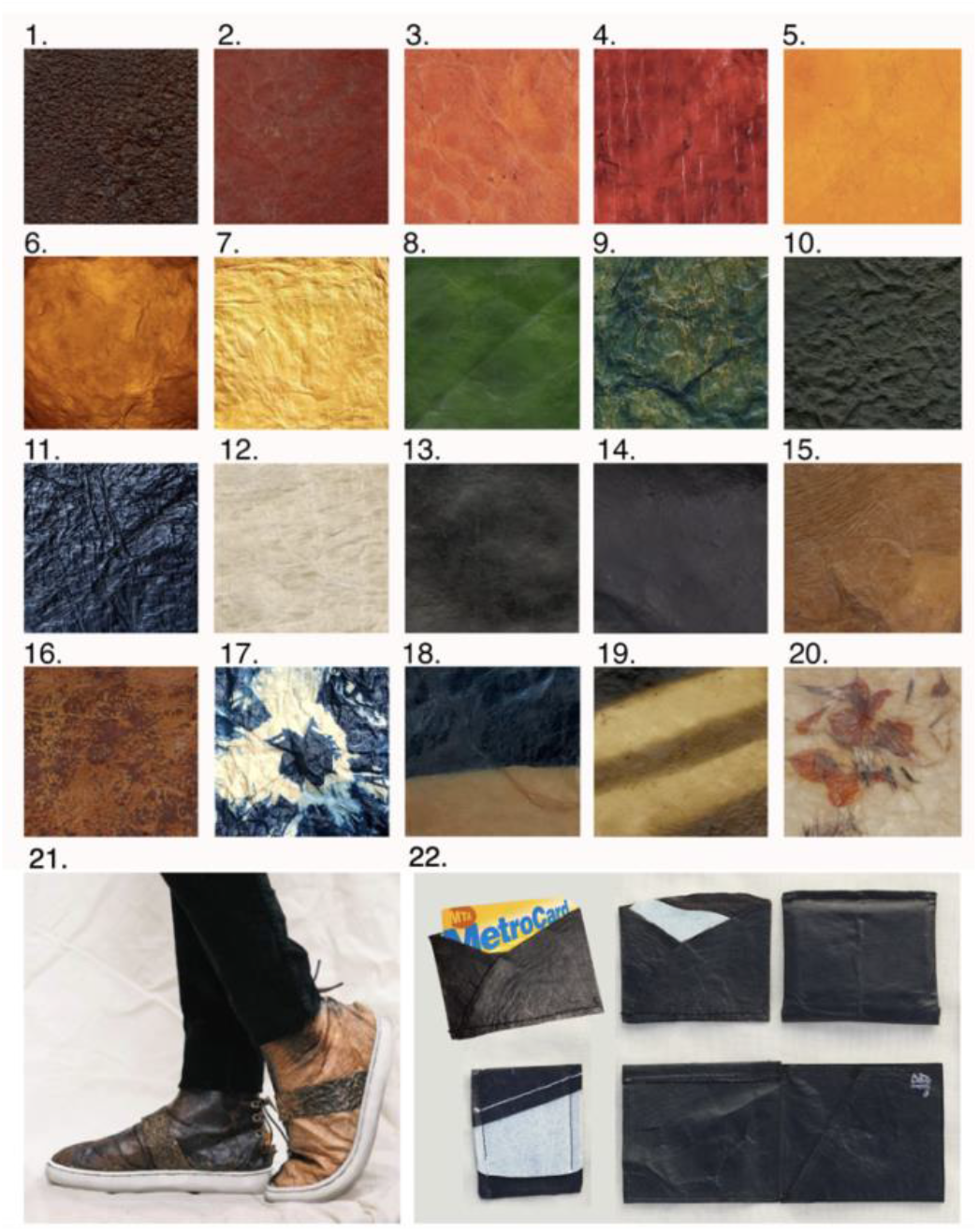
Natural Dye Coloration of Microbial Biotextiles. (1-3) madder root; (4) cochineal, (5) marigold; (6) turmeric; (7) onion skins; (8) chlorophyll; (9-11); indigo after 1,3 and 6 dips sequentially in the dye vat; (12) 0.5 M NaOH; black created through: 13) chemical interaction of tannic acid from black tea and iron acetate, extracted from discarded nails (13); and myrobalan overdyed with logwood and dipped in iron acetate (14); (15-16) as-fabricated MC treated with texture created by application of soy wax heated to 80C. Control of color modulation through: Shibori/Adire folding techniques (17); and partial submergence of biotextile in indigo vat (18). (19) Dark stripes obtained by chemical interaction of iron acetate and tannic acid on as-fabricated MC; and (20) marigold flowers embedded between MC pellicle layers before drying, adapted from Japanese Hapa Zome techniques, showing design opportunities such as no sew seams offered by the layered structure of the moldable biotextile. Design prototypes, including (21) sneakers in natural color with an upcycled rubber sole; and (22) bioleather wallets with natural black as in (14) before lecithin tanning and drying and with white as in (12). (additional prototypes are shown in Fig. 1 and ToC). See Methods for details.

A cradle-to-gate life cycle impact assessment (LCA) is used to compare the environmental and human health impacts of LT biotextile production with the manufacture of conventional textiles, including chrome-tanned leather, synthetic leather (polyurethane (PU)-coated polyester), and woven cotton (Figure 6). For direct comparison, normalization^33^ and weighting factors^34^ are used to convert the manufacturing impacts across eleven categories into a single score in units of milliPoints (mPts)/m^2^ of textile (Table S3); one point (Pt) represents an average individual’s annual share of the total environmental impact in the United States. LT biotextiles biosynthesized in the lab or processed from by-products are denoted biotextile_LAB_ and biotextile_BP_, respectively.

**Figure 6.**
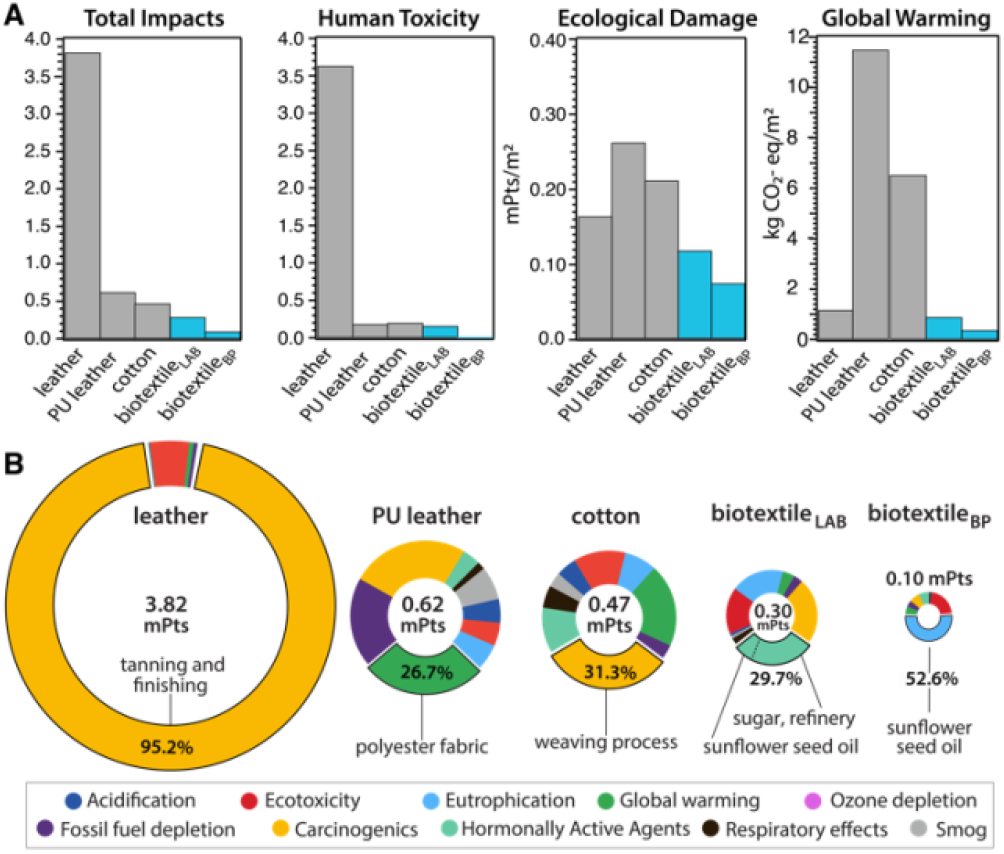
Life Cycle Impact Assessment (LCA). Cradle-to-gate LCA for the industrial manufacture of 1 m^2^ of textile, including cow leather, synthetic leather (PU textile (polyurethane-coated polyester), and woven cotton, compared with microbial biotextiles produced by lab-scale biofabrication (biotextile_LAB_)), and from MC obtained as SCOBY by-product of commercial food and (kombucha) beverage production (biotextile_BP_). (A) Comparison of environmental impact categories for the different textiles: human toxicity corresponds to impacts from carcinogenics and hormonally active agents, and ecological damage includes acidification, ecotoxicity, eutrophication, global warming and ozone depletion. (B) Breakdown of total impact by category, with the largest contribution (material/process) to the impact hotspot (% of total) included.

MC biotextiles offer a significant improvement in environmental performance relative to conventional textiles, especially cow leather, which has the largest total impact of the group (Figure 6A). The dramatic impact reduction relative to leather is overwhelmingly due to elimination of the human toxicity caused by chromium VI derivatives in chrome-tanning, specifically carcinogenics, which account for 95.2% of leather’s total impact (Figure 6A, Table S3). Notably, variants of the lecithin tanning were common pre-Industrial methods of tanning hides, and are still used today, but the process only mechanically softens the hide and requires an additional step to modulate cross-linking in collagen to complete the conversion from hide to leather, usually achieved with an aldehyde (smoke) treatment.^35^

Although two orders of magnitude lower than for cow leather, human toxicity is also the largest impact category for biotextile_LAB_. Refinery processes for sugar, the carbon source in the fermentation media, dominate this toxicity hotspot via carcinogenics and hormonally active agents (Figure 6B). This motivated us to investigate the potential benefit of sourcing MC as a by-product of commercial food and beverage production, in the form of SCOBY pellicles from a local kombucha brewery (biotextile_BP_). After cleaning, lecithin processing and drying, biotextile_BP_ displayed high tensile strength and ductility, comparable to biotextile_LAB_ (Methods, Figure S7).

Elimination of the fermentation media - and the corresponding agricultural and processing impacts— for MC biotextiles yields a 94% reduction in human toxicity (0.01 vs 0.16 mPts) and a 58% reduction in carbon emissions (0.37 vs 0.89 kg CO_2_-eq/m^2^) for biotextile_BP_ compared to biotextile_LAB_ (Figure 6A). Critically, the fermentation media not only accounts for ∼67% of the total impacts (0.1 vs 0.3 mPts) of MC biotextiles, but also 30-65% of the cost of industrial MC production.^36-37^ Thus, the ability to source MC, as well as nutrients for the biosynthesis media, from agro-industrial waste streams opens opportunities for advanced, functional biotextiles with improved circularity and commercial viability.

Biotextile_BP_ mitigates the toxicity of cow leather by a factor of 1000, and the carbon footprint of synthetic leather and cotton by 96.8% and 94.3%, respectively (0.37 vs 11.50 (PU leather) and 6.5 (cotton) kg CO_2_-eq/m^2^). Notably, the Ecoinvent data used to model leather assumes hides are by-products of milk and meat production. As such, the assessment does not include the large climate impacts of the livestock industry,^38^ and the LCA shows a minimal carbon footprint for leather (1.17 kg CO_2_-eq/m^2^) compared to the other textiles (Figure 6). When modeled as a co-product of the livestock industry, with a portion of cattle raising and slaughterhouse inputs and outputs included in the LCA scope, the impacts of leather manufacture increase by two orders of magnitude (128 kg CO_2_-eq/m^2^ and 989 mPts/m^2^ total impact (Methods, Table S4).

Finally, we compare the impacts of in situ coloration of MC biotextiles with yellow onion skins obtained as food by-product/waste, with impacts reported with synthetic dyeing and finishing of cotton. This simple natural process eliminates 348.13 kg TEG ecotoxicity, 0.18 kg C_2_H_3_Cl-eq human toxicity, and 5.4 kg CO_2_-eq of climate impacts from synthetic dyeing and finishing, based on data from Murugesh *et. al*..^39^

Disposal of leather and synthetic leather waste is a serious problem as both are non-biodegradable by microorganisms, while land co-disposal and incineration produce greenhouse gases and introduce carcinogens and other toxins to the ecosystem. In contrast, MC is expected to degrade once exposed to environments rich with cellulolytic microorganisms that hydrolyze the β-1,4-glycosidic linkages of cellulose. To investigate environmental degradability of LT biotextiles, samples (n=5) were weighed and buried at a depth of at least 2.5 cm in tested, nutrient rich soil. After 60 days, with temperatures between 14.2° and 3.2° C, LT showed significant visible deterioration, crumbled easily, and had lost 74.45 ± 2.94 % of the initial mass in the natural environment (Table S8). The data confirms that MC-based products may rapidly reenter the ecosystem, where they may promote the CO_2_ capture potential of healthy soil, and support a closed loop material life cycle.

## Conclusions

Harnessing biofabrication and inspired by ancient practices, we engineer high-performance microbial nanocellulose bioleather with a sustainable circular life cycle. This biotextile approach eliminates the use of hazardous substances such as chromium VI derivatives for tanning, halogen compounds for fire-retardancy, petrochemical feedstocks and phthalates as plasticizers for performance properties, and synthetic dyes for color, which make the textile industry a leading source of global waste water and microplastic pollution. Life cycle assessment reveals an order of magnitude reduction in total environmental damage for production of lecithin-tanned biotextiles relative to conventional textile manufacture, and highlights how waste-to-resource strategies can further mitigate impacts. In particular, the microbial bioleather offers a thousandfold reduction in toxicity of chrome-tanned leather and an order of magnitude lower carbon footprint than synthetic leather and cotton.

Future prospects include MC-based alternatives to cotton and synthetic fibers as well as conductive, biomedical and sensor textiles, with functional properties directed programmed into microbial biosynthesis, and further tailored by environmentally benign processing.^36^ Beyond textiles, phosphorylated biopolymers, including MC, show high performance for diverse applications, from heavy metal pollution remediation to tissue culture and bone regeneration,^40^ but the many potential benefits are offset by the toxicity and non-renewability of reagents and high temperatures of conventional phosphorylation processes. The process developed here may therefore inform efficient green manufacturing routes for biopolymers to realize their full potential in a regenerative bioeconomy. Thus, advances in microbial cellulose performance incentivizes a transition to a circular materials economy, characterized by rapidly renewable resources, low impact processes, scalability, and biodegradability.

## Author Contributions

Conceptualization: TNS, RA, HHL. Methodology: TNS, RA, AMVC, SG, AC, HHL. Investigation: TNS, RA, STR, DA, SE, Y-TC, CTJ. Visualization: TNS, RA, DF, HHL. Funding acquisition: TNS, HHL. Supervision: TNS, HHL, SG, AMVC, CN, SKK. Writing – original draft: TNS, RA, DF, HHL, Writing – review and editing: All authors.

## Acknowledgments

We thank OM Champagne Tea (Mount Kisco, NY) for donations of microbial nanocellulose, produced as a fermentation by-product of food and kombucha beverage production. The bioleather sneakers shown in the ToC figure were designed by and created in collaboration with Dao-Yi Chow and Maxwell Osborne of Public School, for the OnexOne Conscious Design Initiative. This work was supported by the National Science Foundation, MRSEC program through Columbia on Precision-Assembled Quantum Materials (PAQM), grant DMR-2011738 (TNS, RA, Y-TC, HHL, CTJ, SE, DA, STR), the National Institutes of Health, grant NIH-NIAMS 1R01-AR07352901 (HHL), and a DoD CDMRP award, grant W81XWH-15-1-0685 (HHL).

## Data and Materials

All data are available in the main text or the supporting information.

## ToC

Harnessing microbial biofabrication coupled to a processing protocol inspired by indigenous textile processes, we engineer high-performance biotextiles with a sustainable circular life cycle, including the plant and mineral dyed bioleather sneakers shown.

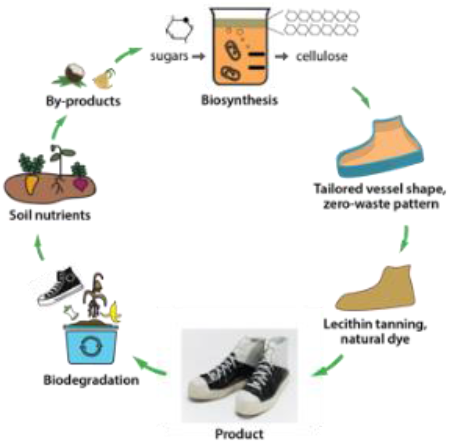

