## Supplementary Materials for "Microbial Nanocellulose Biotextiles for a Circular Materials Economy"

**This PDF file includes:**

Materials and Methods
Supporting Information
Figs. S1 to S7
Tables S1 to S9
Captions for Movie S1 and S2

**Other Supporting Information for this manuscript include the following:**

Movie s1
Movie s2

### Materials and Methods

#### Biosynthesis of microbial cellulose pellicles

Microbial cellulose pellicles were prepared in culture media containing 5.8 w/v% sucrose as a carbon source, 2% w/v green tea as a nitrogen source, and Symbiotic Colony of Bacteria and Yeast (SCOBY) obtained from Fermentaholics © or provided by OM Champagne Tea, a commercial kombucha beverage facility. Culture media was inoculated with 10 w/v% SCOBY starter culture consisting of a combination of bacteria and yeast. Static fermentation at room temperature was maintained until a pellicle at least 2 cm thick was formed at the air-liquid media interface. Biofabrication proceeds as 2-D layer by layer production of a cellulose biofilm which takes the shape of the fermentation vessel. Formed pellicles were washed three times with distilled water to remove residual sugars and air dried on a polypropylene tray at room temperature, generally 48-72 hours for the 1-2 inch pellicles used in this study.

As a post-production treatment, hydrated pellicles were rinsed three times in distilled water, and directly immersed in a lecithin emulsion for 24-48 hours (LT MC). The lecithin emulsion (LT) consisted of 5% w/v lecithin powder (soy or sunflower seed, 97% phosphatidylcholine) created as a byproduct of the edible oil industry, and 20% v/v sunflower seed oil in water and blended at high speed for 60 seconds to create an emulsion. After removal from the emulsion, LT MC samples were rinsed three times in distilled water and air dried on a polypropylene tray at room temperature under the same conditions as the as-fabricated MC control. Smoke treatment was performed on dried (as-fabricated or lecithin tanned) samples.

Microbial nanocellulose samples were also processed following traditional brain tanning practices; the increased mechanical performance for brained tanned MC provided a positive control to investigate the effect of lecithin tanning. A calf brain was purchased from Esposito Meat Market, New York, NY, and processed into an emulsion with a blender. The brain emulsion was rubbed into the MC surface, left for two hours, and then rinsed.

Aldehyde smoke treatment was performed on dried samples. Dehydrated MC samples, both as fabricated and lecithin tanned, were exposed to an additional smoke tanning aldehyde treatment, in which biofilms were placed into a hydrocarbon-rich environment for 1 hour, using a Smoke Hollow 26142E 26-Inch Electric Smoker with Adjustable Temperature Control and Kingsford smoking wood chips, with the temperature between 160° and 210°F. Prolonged exposure of the MC biotextiles to hydrocarbon-rich smoke was investigated as a mean to seal in the lecithin and oils, which has been documented to traditionally follow braining to complete the tanning process.

Characterization

Microbial cellulose morphology was assessed by using a scanning electron microscope (Zeiss Sigma VP, Oberkochen, Germany; 3 kV; n=5). Briefly, hydrated microbial cellulose samples were initially placed in a -20$^{o}$C freezer for 24 hours and lyophilized in a freeze dryer system (Labconco FreeZone, Kansas City, MO, USA) for 24 hours at −84 °C and 2.0 × $10^{-2}$ mbar. Prior to imaging, samples were sputter coated (Cressington 108, Watford, UK) with 30 nm of gold. The fiber diameter of each sample was measured by analyzing randomly selected fiber segments in SEM images using NIH ImageJ software (Bethesda, MD, USA; n=100), to calculate an average fiber diameter.

Biotextile hydrophilicity was measured via water contact angle (WCA), in which angles less than 90 degrees are indicative of hydrophilic surfaces, while greater than 90 degrees are characteristic of hydrophobic properties. Room temperature deionized water (10 uL) was deposited onto microbial cellulose surfaces, and contact angles were measured using a contact angle goniometer (Ramé-hart, Inc., Mountain Lakes, NJ, USA) with a fiber optic illuminator (Dolan-Jenner Industries, Inc., Woburn, MA, USA).

Mechanical properties of microbial cellulose biotextiles were assessed by securing samples with custom clamps and mounting in a uniaxial tensile testing machine (Instron, Model 1321, Norwood, MA, USA), equipped with a 25 kN load cell. Biotextiles were maintained to have a gauge length of 2 inches and were tested to failure. Microbial cellulose elastic modulus, toughness, and ultimate tensile strength were determined from the stress-strain curve.

X-ray Diffraction

A 5-10 mm punch pressed into each of the various treated samples to create a series of small disks. The disks were placed on a standard glass slide. Diffractograms were collected using a Panalytical XPert3 Powder diffractometer with a 3 kW generator, fully ceramic Cu Long fine focus (LFF), X-ray tube, vertical goniometer (theta-theta) and a PIXcel 1d detector. The copper long fine focus lens produces a K- alpha of 1.5406 angstroms. Measurements were collected while scanning from 5 to 100 degrees with the following parameters: Fixed divergence slit: 1/2 degree, step size: 0.026 deg/step, scanning rate: 10.7 deg/min. A background was obtained by measuring the blank sample holder before taking measurements. A baseline correction was applied using Highscore XRD software.

The lattice spacing (d-spacing) was calculated using Bragg's equation:

$$\lambda = 2d_{hkl} * \sin\theta$$

where $d_{hkl}$ is the lattice spacing of the crystallographic planes, θ is the corresponding Bragg angle, and λ is the X-ray wavelength (0.154 nm).

The Crystallinity Index (CI) was calculated using the XRD peak deconvolution method[1] using Igor-Pro software to separate amorphous and crystalline contributions to the diffraction spectrum using a curve-fitting process. For the curve fitting, Gaussian functions were used for deconvolution of XRD spectra into four crystalline peaks (101, $10\bar{1}$, 021, and 200) and one amorphous peak centered at a 2θ value of ~21.5 degrees. CI is calculated from the ratio of the area of all crystalline peaks to the total area.

It is generally accepted that peak broadening is due to the amorphous cellulose, but crystallite size is an equally important issue for peak broadening and can affect accuracy of CI values obtained with the deconvolution method. However, uniformity in average crystallite size for as-fabricated MC and processed (LT, LTS, S) biotextiles, calculated from this method using the Scherrer formula, validate the accuracy of using this approach, and attributing differences in Bragg peak ratios to amorphous and crystalline contributions (see **Table S1**).

Crystallite Size was obtained by calculating the Scherrer width via the Scherrer equation:

$d_{hkl} = k\lambda/(FWHM * \cos\theta)$

where $d_{hkl}$ is the size of the crystallite in nm, λ is the x-ray wavelength, is the FWHM of the peak, k is the Scherrer constant of 0.94 and θ is the diffraction angle (2θ is the corresponding peak position).

Thermal Analysis

A 45º Flame Test (ASTM D1230-94), which is a widely-accepted method for determining apparel textiles flammability, was used to measure: 1) Ease of textile ignition, and the 2) Duration of flame spreading. Biotextiles (1"x 6" in **Fig. 3** and Fig. S2 and 1"x18" in **Fig. S3)** were mounted and held in a custom apparatus at an 45º angle. A standardized flame was applied to one end of the biotextile at a 45º angle, using 14.1 oz. Map-Pro Cylinder by Bernzomatic, which uses liquid propane as a fuel source and has a flame temperature in air of 3,730° F/2054.4° C.

Charred biotextile surfaces were observed with SEM. Thermal properties and char formation of dried, pre-weighed microbial cellulose discs (diameter: 4.5 mm) were determined by a thermogravimetric analyzer (TGA 550, TA Instruments; New Castle, DE). TGA analysis was performed at a heating rate of $10^{o}C/min$ over the temperature range of $25^{o}C$ to $700^{o}C$ under flowing nitrogen (40 mL/min), in which final weights indicated the formation of char residue.

Chemical and elemental analysis via FTIR, XPS, and EDAX were performed to characterize microbial cellulose mechanical, hydrophilic, and thermal properties. FTIR (LUMOS II, Bruker, Billerica, MA, USA) spectra were recorded under attenuated total reflectance (ATR) mode in a spectral range of 4000-600 $cm^{-1}$. Each spectrum was collected using a total of 200 scans and a resolution of 4 $cm^{-1}$.

Cellulose chains contain both crystalline (ordered) and amorphous (unordered) regions. To determine the relative amount of crystallinity in materials, the Crystallinity Index (CI) can be determined by XRD, $^{13}C$ and FTIR.

Determination of CI from FTIR has been found to be the easiest, however, these results only provide relative values because cellulose contains both crystalline and amorphous regions.

Crystallinity analysis of the biotextiles with FTIR data was performed using empirical formulas proposed by Nelson and O'Connor.[2] The first calculation is the Lateral Order Index (LOI: $A_{1420}/A_{893}$), also known as the empirical crystallinity index, relates to the overall degree of order in cellulose. The peak around 1420 cm$^{-1}$ relates to the presence of crystalline regions, while the peak around 893 cm$^{-1}$ is associated with amorphous regions. The second calculation is the Total Crystallinity Index (TCI: $A_{1375}/A_{2900}$) is also used to evaluate crystallinity. While the ratio between absorbances at 1429 cm$^{-1}$ to 897 cm$^{-1}$ is defined as the lateral order index (LOI) required to assess cellulose's overall degree of order, the ratio between 1372 cm$^{-1}$ to 2900 cm$^{-1}$ is defined as total crystallinity index (TCI), which is related to the degree of crystallinity in cellulose. The formulas used for these calculations are as follows:

$$\text{LOI} = \frac{A\,(1420)}{A(\,893)}$$

$$\text{TCI} = \frac{A\,(1375)}{A(\,2900)}$$

<u>Determination of Hydrogen Bonding Environment Via FTIR</u>

The relative absorbance (A) for the band at a specific wavelength (λ) attribute to the hydrogen bonds in the material is calculated using the C–H stretching band as an internal standard. The empirical formula is:

$$Ar_{\lambda} = \frac{A_{\lambda}}{A_{C-H}} = \frac{A_{897,\,1375,1430}}{A_{2920}}$$

The bands 2920 cm$^{-1}$, 1372 cm$^{-1}$ and 1423 cm$^{-1}$ are assigned to the C–H stretching in the methylene groups, aliphatic C–H stretching, and aromatic skeletal vibrations combined with C–H in plane deformations, respectively. According to the empirical formula, the absorbance ratio of the bands at 897 cm$^{-1}$ ($Ar_{897}$), 1375 cm$^{-1}$ ($Ar_{1375}$), and 1430 cm$^{-1}$ ($Ar_{1430}$) to 2915 cm$^{-1}$ correspond to the degree of hydrogen bonds destroyed. The rule of thumb is a decrease in ratio suggests that there is an increase in hydrogen bonding strength (*ref. (31) in main text)*.

Molecular level details of surface bonding of lecithin tanned and as fabricated MC biotextiles were determined by XPS (PHI 5500, Chanhassen, MN, USA) after samples were vacuum annealed at 75°C 24 hours. The spectra were recorded using a monochromatic Mg-Ka radiation X-ray source (1253.6 eV) and the analyzer pass energy was set to 25 eV. The sample chamber was set to 50 W operating at 15 kV voltage and a base pressure of 2 x10$^{8}$ torr. The XPS spectra were collected in the range from 0 to 1200 eV, with a resolution of 0.1-1.0 eV. A Shirley background subtracted was performed to remove the inelastic background of the carbon (C 1s), oxygen (O 1s), nitrogen (N 1s), and phosphorous (P 2p) electron core spectra and data was analyzed using commercial curve fitting software Igor64 (WaveMetrics, Portland, OR, USA). The binding energy scale was calibrated using the Au 4f 4/7 line of 10 nm thick gold, electron beam evaporated into 5 mm wide strips onto two parallel edges of the LT and MC samples, and confirmed against the C 1s binding energy for adventitious carbon on Au (285 eV) as well as published XPS data for microbial and plant cellulose. Based on these values, MC and LT XPS data were shifted -1.98 and -2.88 eV to higher binding energy, respectively. A smoothing factor of 10 was applied to the P 2p XPS data for ease of comparison (no smoothing was applied to C 1s or O 1s data).

Further surface elemental characterization was assessed by an EDS system (Bruker XFlash 6, Billerica, MA, USA; 10 kV; 5 minutes; n=3) connected to a scanning electron microscope (Zeiss Sigma VP, Oberkochen, Germany). Biotextiles were sputter coated (Cressington 108, Watford, UK) with copper, and assessed for the presence of phosphorus.

<u>Natural (plant, insect and mineral) dyeing of microbial cellulose biotextiles</u>

To reduce the water and energy demands of traditional dip dye methods, natural dye matter was added with tea and sugar for coincident extraction of color and preparation of the cultivation media and strained before adding bacteria inoculum to the media- so that color could be directly incorporated into the nanofiber mesh during biosynthesis. The tannins in the tea serve to mordant the cellulose fibers, to improve color retention, saturation and colorfastness. Cultivation vessels were prepared with 5.3% w/v sugar, 4% w/v tea, 10% v/v starter culture and natural dye matter.

Dye procedures are as follows. A madder root 9.75% w/v post synthesis dye bath was used for **Figure 5.1**. The following dye stuff was incorporated directly into the culture media so that color was incorporated into the microbial cellulose matrix *during* biosynthesis: 10% w/v madder using black tea (**Figure 5.2)** and green tea **(Figure 5.3**) as the nitrogen source in the fermentation media (the swatch shown in Figure 5.3 was dried from a pellicle grown to half the hydrated state thickness of that in **Figure 5.3**); 7.5% w/v cochineal (**Figure 5.4**); 5% w/v; marigold extract (**Figure 5.5**); turmeric 20% w/v; (**Figure 5.6**); 7.5% v/v liquid chlorophyll (**Figure 5.8**). The swatch in **Figure 5.7** was colored using yellow onion skins 3% w/v as a post processing dye bath, after color was removed using the process described below for the white swatch shown in **Figure 5.12**. Onion skins were obtained as food by-product and chlorophyll was obtained from Horbȧach, Melville, NY. Marigold flowers were harvested locally at the end of season bloom and embedded between layers of the hydrated MC biofilm and hammered flat using the ancient Japanese textile art technique of hapa-zome, before the biotextile was left to dry flat on a flat polypropylene tray under ambient conditions (**Figure 5.20**). All other dye stuff was obtained from Botanical Colors, Seattle, Washington.

For red shown in main text **Figure 5.4** and **Figure 1**, 10 grams of cochineal were ground to a fine powder, placed in a cotton muslin pouch and tied with cotton string, and immersed in 250 ml distilled water and brought to 100 ºC for 15 minutes. The heat was turned off, allowed to sit for 15 minutes, after which the process of adding water and bringing to a boil was repeated 3 times. On the last step the tea and sugar for the culture media were added, after which the cochineal pouch was removed, the additional water for the culture media was added, cooling the media to a temperature suitable to add the inoculum (10% v/v). Other colors were extracted in one liter of water held at 85ºC to 45-1 hour, and either added directly to the biosynthesis media after cooling or used as a post-processing dip dye bath, except for chlorophyll, which was added at room temperature with the inoculum to the media.

Post synthesis dip dyeing in room temperature organic indigo vats, produced from combining 5 g organic indigo powder, 10 g calcium hydroxide and 15 g fructose in 15 liters of water were used to create greens (**Figure 5.9**) and blues (**Figure 5.10-5.11, 5.17**). White biotextiles were achieved by immersion of hydrated MC in 0.5 M NaOH at room temperature for 12 hours, followed by rinsing in distilled water until a neutral pH was reached (**Figure 5.12**). Black coloration on the biotextile shown in **Figure 5.13** was achieved by chemical reaction of tannic acid and Fe acetate. Samples were immersed in a 3% w/v tannic acid solution, obtained from Botanical Colors, followed by a dip in aqueous Fe acetate mordant, prepared by soaking 100 grams of nails in 2 liters of 2.5% v/v acetic acid for two weeks. Black biotextiles shown in **Figure 5.14** and used for the prototypes shown in **Figure 5.22** were created in a post-processing overdye procedure as follows: 10% w/v myrobalan bath, brought up to 55ºC before MC biotextiles were added, and heating was continued to a high simmer (~83ºC) and held for one hour. MC biotextiles were removed from the myrobalan bath, rinsed twice, and left to cool before being added to a 15% w/v logwood extract bath at 33ºC. The logwood bath was held at this temperature for 30 minutes, after which the temperature was gradually increased to 80ºC) for 45 minutes. MC biotextiles were rotated gently throughout the post-synthesis dye process, and left to cool in the dye bath, then rinsed twice before being dipped in Fe acetate for 10 seconds, and rinsed three times.

All post-processing dye baths were saved for repeated use (usually three more dye processes) until color diminished. After harvesting colored MC pellicles, all in situ dye baths were resupplied with cultivation media nutrients for at least 3 harvests.

Variation in surface texture shown in **Figure 5.1, 5.15 and 5.16** was achieved using soy wax heated to ~83ºC, spread across the biotextile surface with a brush, and pressed against an aluminum foil-covered hot plate for 10 seconds. Other surface variation was introduced by drying with tension (edges of biotextile clipped to a flat surface, **Figure 5.13-5.14; 5.18-5.19**) or without tension, as shown in **Figure 5.7-5.11**, or dried on a screen (**Figure 5.4**).

Life cycle impact assessment (LCA)

LCA analysis is a quantitative technique to assess the environmental and human health impacts associated with all the stages of a product's life, which is from raw material extraction through materials processing, manufacture, distribution, and use as specified in ISO 14040 standards, including International Organization for Standardization (ISO). Environmental Management - Life-Cycle Assessment-Principles and Framework, International Standard 14040; ISO: Geneva, Switzerland, 2006. International Organization for Standardization (ISO). Environmental Management - Life Cycle Assessment - Requirements and Guidelines, International Standard 14044; ISO: Geneva, Switzerland, 2006.[3] Manufacturing impacts were determined using Sustainable Minds® Life Cycle Assessment software (Cambridge, MA, USA) and the EcoInvent database (Zurich, Switzerland). The impact categories are based on the U.S. EPA's Tool for the Reduction and Assessment of Chemical and other Environmental Impacts (TRACI) life cycle impact assessment (LCIA) methodology. As defined by the software, ecological damage is comprised of acidification (kg $SO_2$ eq), ecotoxicity (CTUe), eutrophication (kg N eq), global warming (kg $CO_2$ eq), and ozone depletion (kg $CFC^{-11}$ eq). Human health damage includes carcinogenics and non-carcinogenics (both in units of CTUh), respiratory effects (kg PM2.5 eq, fine particulates), and smog (kg (ground level) $O_3$ eq), while resource depletion exclusively refers to extraction of fossil fuels (MJ surplus).

Impacts were calculated in a cradle-to-gate LCA comparing the impacts of biofabrication microbial biotextiles using the methods described above with the manufacturing impacts of conventional textiles, including chrome tanned leather, synthetic leather (polyurethane (PU)- coated textile), and woven cotton. The functional unit is $1m^2$ of textile. For direct comparison, impacts were calculated from weighted impact categories, in milliPoints(mPts)/$m^2$) of textile, utilizing normalization and weighting factors shown in **Table S3**.[4] A point (1000 mPts) represents the average person's annual environmental load (*i.e.* entire production/consumption activities in the economy) in the United States.[5] Normalization factors were calculated using characterization factors from the TRACI 2.1 LCIA model, and toxicity-based categories use characterization factors calculated with USEtox. Weighting factors represent degree of immediate concern or degree to which remedial actions associated with the impact are underway, and provide a practical method to link quantitative results of LCA with environmental performance of competing products and assist environmentally preferable design, manufacture, and purchasing.[6] A cradle-to-gate partial product life cycle was performed from resource extraction (cradle) to the factory gate which includes raw material and manufacturing impacts, with distribution, consumer use and disposal omitted; such assessments are often used as the basis for environmental product declarations (EPD).

LCA Rationale: Market data of the leather and synthetic leather goods industry.
Both synthetic and animal leather are frequently utilized within the apparel, automotive, and furnishing industries. According to Grand View Research.[7] The global synthetic leather market size was valued at 29.3 billion USD in 2019 and was expected to increase to 30.3 billion USD for 2020. It is also estimated that the market size volume in 2020 accounted for 15,585.6 million meters. The market size value of animal leather is estimated to have been over USD 80 billion in 2019.[8] As reported by Grand View Research, the textile market size value in 2020 is USD 1,000.30 billion. Subsequently, cotton was anticipated to be the largest raw material segment, accounting for a market share of 39.5% in 2019.[9]

Synthetic Leather

Synthetic leather (polymer coated textile) can be produced with a variety of synthetic polymers, including polyvinyl chloride (PVC), polyvinylidene chloride (PVDC), polyurethane (PU), and different base fibers (cotton, polyester or cotton/synthetic blends), and coating techniques. Each one of these variables affects the ecological impact of synthetic leather. The variation chosen for this LCA is PU-coated polyester made by the company Kuraray, specifically their product called Clarino Crust. According to market research, this enterprise is one of the main manufacturers of synthetic leather, and the material inventory created in the EcoInvent database are based on the manufacturing processes used by Kuraray.[10] According to Kuraray's Clarino Crust technical sheet, the weight of the textile is 0.66± 0.02 kg/m$^2$ and contains 90% of polyester (PET)- 0.594 kg - and 10% polyurethane (PU) -0.066 kg. The only information taken from the Clarino Crust technical sheet is the weight and the percentage -of PU and PET- utilized.[11]

The polyester was modeled with average data for dyed, manufactured polyester fabric for apparel in the EcoInvent database within the Sustainable Minds software. The PU coating system of materils was built from the average values material inputs for the urethane composition of use in synthetic leather.[12] In particular, *t*he urethane composition is the dry method urethane composition. In parts by weight, the dry method urethane composition includes lower components: Solvent type polyurethane resin 80-100 parts (average 90 parts), acid additive 1-2 parts (average 1.5 parts), dimethylformamide 5-30 parts (average 17.5 parts) and ethyl acetate 10-20 parts (average 15 parts). *T*he inventory included 59.4 g epoxy (PU) resin (CAS number: 025928-94-3), 9.9 g ethyl acetate (CAS number: 000123-86-4), 1 g citric acid from fermentation (1 g), 11.55 g N,N-diemthylformamide (CAS number: 000068-12-2), and polyester fabric (594 g). Electricity, water use and transportation were not included.

For comparison, we also modeled the material inputs for the PU coating using the data set for PU flexible foam included in the EcoInvent database within Sustainable Minds, CAS number: 009009-54-5, provided by Kuraray, which includes the transport of the monomers, the production (energy, air emissions) of the PU foam, and using average values of transport and infrastructure, for the typical composition for European conditions and present technology used in Europe. The two models for the PU coating yielded similar impacts values and distributions across categories. The process of foaming by expanding plastic involves converting the plastic foam into an emulsion and then coating the base fabric.[13] The process data set included the auxiliaries and energy demand for the conversion process of plastics and the converted mass of plastic was not included in the dataset. The complete system of materials used to model the manufacturing of PU-coated polyester, obtained from the Ecoinvent database, is shown in **Tables S4** and **S5.**

Cow leather

The functional unit is 1m$^2$, corresponding to a measured mass of 0.8 kg based on measurement.

*Leather as a by-product:* Material inputs for the manufacture of cow leather tanned with a chromium process from the Ecoinvent database and shown in **Table S4.** According to the database, the inventory assumes the hides are a byproduct of milk and meat production without significant value, and therefore neglects the inputs of 989 kg of food and 0.56 ha of grassland per cow, based on average data from the Netherlands. Since less than 10% of revenue of cattle comes from by-products, with leather accounting for the majority of this revenue, a by-product model is compared in Figure 6.[14]

*Leather as a co-product*: This inventory starts with the production of hides and skins as a co-product from beef cattle at a slaughterhouse. The dataset includes (portions of) the following process: cattle raising, slaughtering, preservation of hides, chrome tanning, and finishing of chrome tanned leather. The allocation model used for the production of hides is economic allocation for the slaughterhouse. The inputs and outputs for subsequent processes, which includes preservation of hides, tanning, and finishing is based on Joseph and Nithya, 2008.[15]

The majority of chrome tanning occurs in India, so energy processes for India have been used. Synthetic tans used for tanning is assumed to be a combination of DHDPS and formaldehyde.

Woven Cotton

Data from US-EcoInvent 2.0 is used to model the manufacture of 1 $m^2$ of medium weight (250 $g/m^2$) woven cotton farmed in the U.S., and includes: cotton fiber production, yarn production and weaving processes. The inventory includes the processes of soil cultivation, sowing, weed control, fertilization, pest and pathogen control, irrigation, harvest and ginning. Machine infrastructure and a shed for machine sheltering is included. Inputs of fertilizers, pesticides and seed as well as their transports to the farm, and the direct emissions on the field are also included. The system boundary is at the farm gate. Inputs are based on average production in the USA based on data collected in the period 2001-2006.Water for irrigation is pumped from 48 meter depth by electric pumps. Production of yarn from ginned fibers and weaving of cotton yarn into a textile are also included. The manufacturing impacts are shown in **Table S4**.

Microbial Cellulose biotextiles

*Biotextile$_{LAB}$:*

The system bill of materials was created from the Ecoinvent data sets and shown in **Table S6**. The biosynthesis cultivation media to biofabricate 1$m^2$ of microbial cellulose included 1.74 kg of sugar, 120 grams of green tea, 30 liters of water and 3864.68 Btu of heat energy from a natural gas boiler needed to raise the temperature of 15 liters of water from 25 C to 90 C ($Q=mc\Delta T$) to steep the tea and dissolve the sugar used in the cultivation media. Half the total water used in the liquid media (15 liters) is heated, the other half is added at room temperature to cool the media to a sufficiently low temperature (38 C or less) before adding 10% v/v starter culture to the media. The Ecoinvent data set for sugar included production of sugar from sugarcane and manufacturing at the sugar refinery. Impacts for production of 120 grams of tea used to produce 1 $m^2$ microbial cellulose were calculated based on APOS, U (of project Ecoinvent 3 {LK}) data for 1 kg tea, which includes farm cultivation, including irrigation, land use, field spraying insecticides and fertilizers (ammonium nitrate, ammonium sulfate, glyphosate, diammonium phosphate production, drying and processing. Material inputs for the lecithin tanning emulsion (LT) included 0.28 kg of sunflower oil and 2 liters of distilled water to prepare the emulsion, and another liter of water to rinse after tanning. The sunflower seed oil data set included sunflower seed production and oil mill process. Phosphatidylcholine (lecithin powder) is produced as a byproduct from the edible oil industry, and as such, impacts are considered accounted for in the data set for sunflower seed oil, and is not available as a separate material input.

*Biotextile$_{BP}$: Purification and Processing (LT) of 1 $m^2$ of kombucha fermentation by-product (symbiotic colony of bacteria and yeast (SCOBY)).*

The system of materials was based on the Ecoinvent data set and shown in **Table S7**. In the purification process of the MC in the form of a symbiotic colony of bacteria and yeast (SCOBY) provided by OM Champagne Tea, 8 liters of water and the heat energy to rise to 85C were utilized. The material inputs for the lecithin tanning process included 200 ml sunflower seed oil and 2 liters of water to tan 1$m^2$ of microbial nanocellulose biotextile.

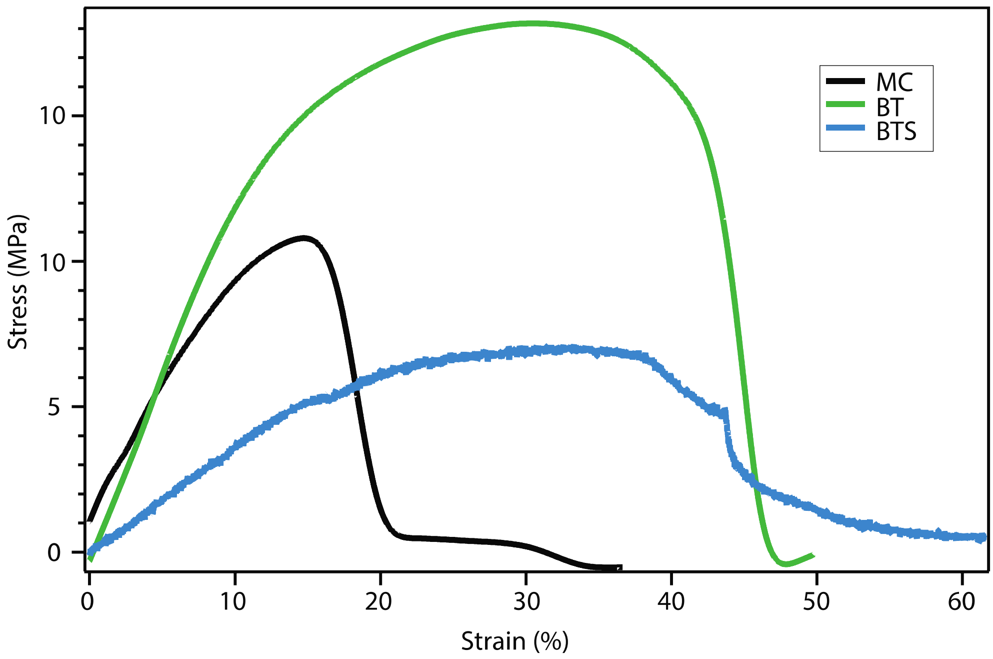


**Figure S1**

Stress-strain curves for brain tanned (BT) and brain tanned and smoked (BTS) microbial nanocellulose (MC), following ancient tanning practices, compared to as fabricated MC control, which encouraged the line of investigation in this work.

| | Element Mass (%, n=3) | |
|---|---|---|
| Element | MC | LT |
| Carbon | 74.68 ± 2.76 | 75.82 ± 1.91 |
| Oxygen | 21.20 ± 2.60 | 19.73 ± 1.97 |
| Nitrogen | 3.00 ± 1.16 | 0.56 ± 0.26 |
| Phosphorus | 0.57 ± 0.18 | 2.02 ± 0.18 |
| Potassium | 0.26 ± 0.05 | 1.71 ± 0.12 |
| Sulfur | 0.25 ± 0.07 | 0.12 ± 0.03 |
| Copper | 0 | 0 |
| Sum | 100 | 100 |

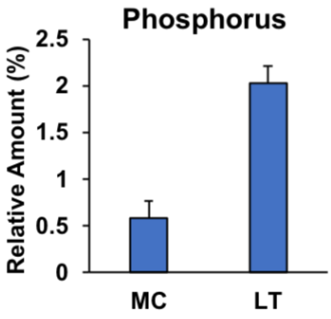


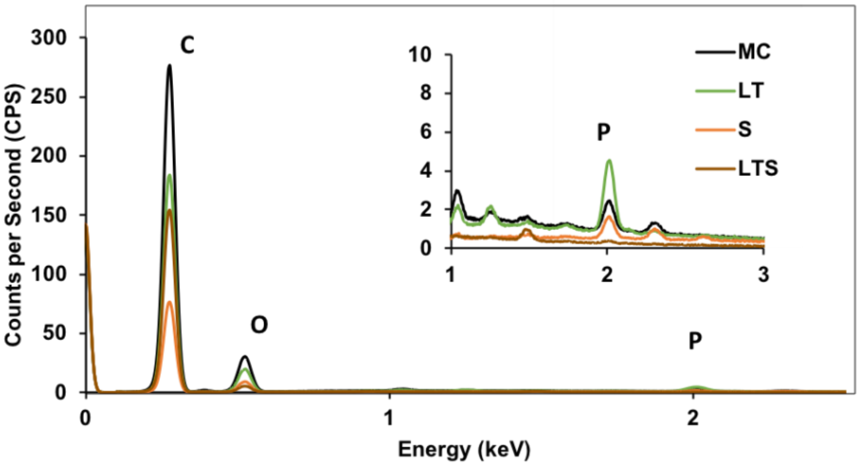


**Figure S2**
Energy dispersive x-ray measurements (^p<0.05, n=3) showing variation in chemical composition for the SEM micrographs in Figure 2A.

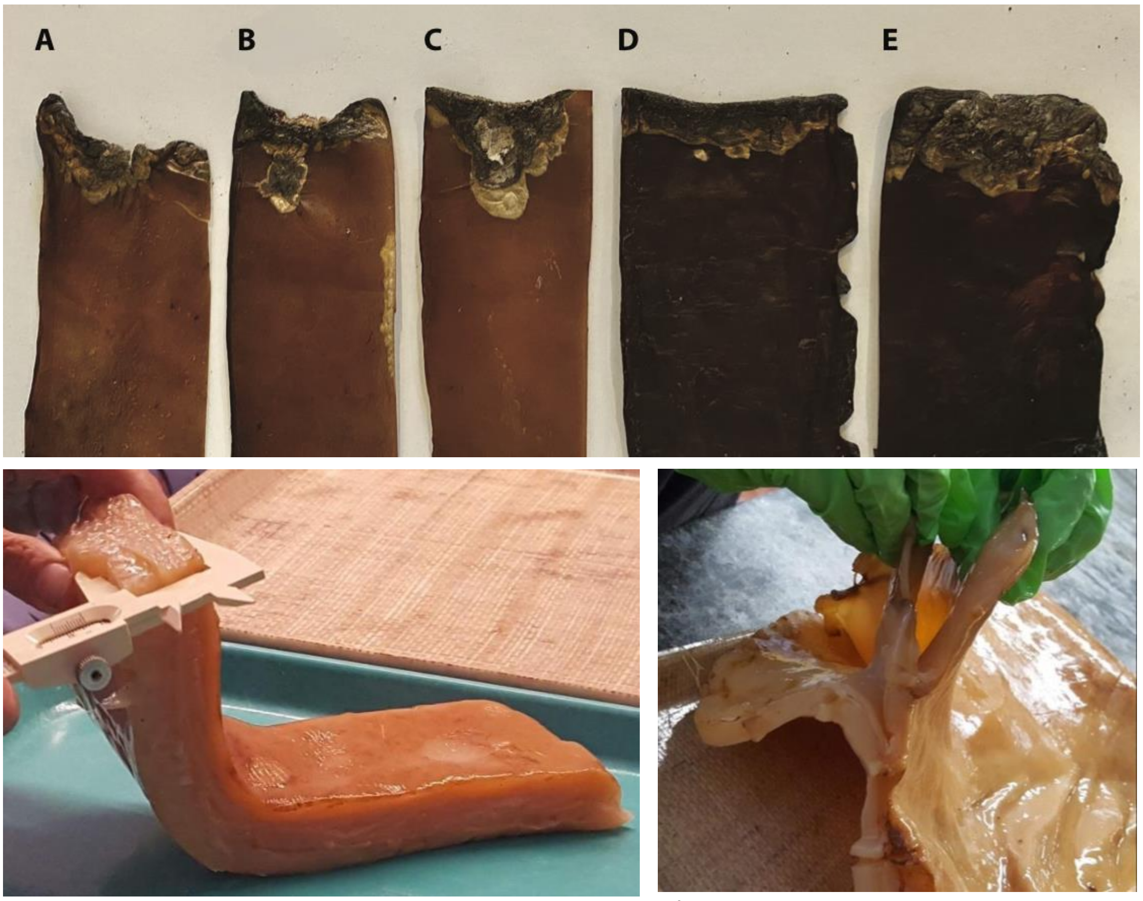


**Figure S3**

**Top**: Flame resistance as a function of MC biofilm thickness obtained from a single pellicle separated into layers with hydrated (pellicle), dehydrated (biotextile) thickness: **A)** (3, 0.15 mm); **B)** (7, 0.26) mm; **C)** (10, 0.41) mm; **D)** (22, 0.71) mm; and **E)** (2.5, 0.90) mm. **Bottom**: Sections of hydrated pellicle before dehydration used for flame study as a function of MC thickness (**left**: (E) 25 mm thickness, **right:** constituent layers **(**A-C, 0.5-0.41 mm**)**.

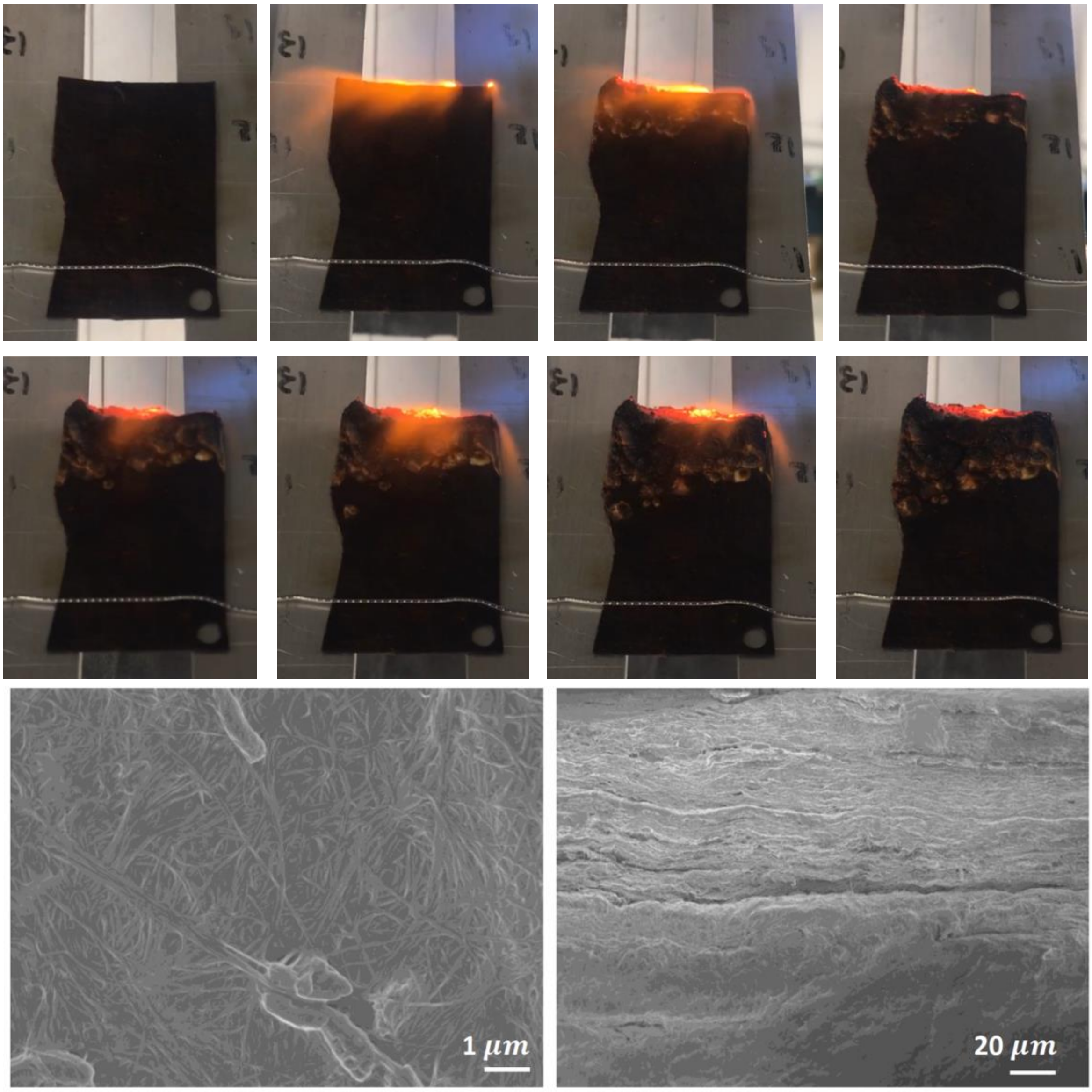


**Figure S4**

**A)** Flame testing of a cast slurry of MC, creating by blending MC pellicles in a Vitamix 3 HP blender and casting a 1.6 cm layer on a screen to dry in under ambient conditions. **B)** SEM images of untreated MC slurry surface **(left)** and cross-section **(right)** showing the blended MC self-organized into a layered structure, which as for the as grown MC films, imbues the material with a degree of flame retardance unique to microbial cellulose (as compared to plant cellulose).

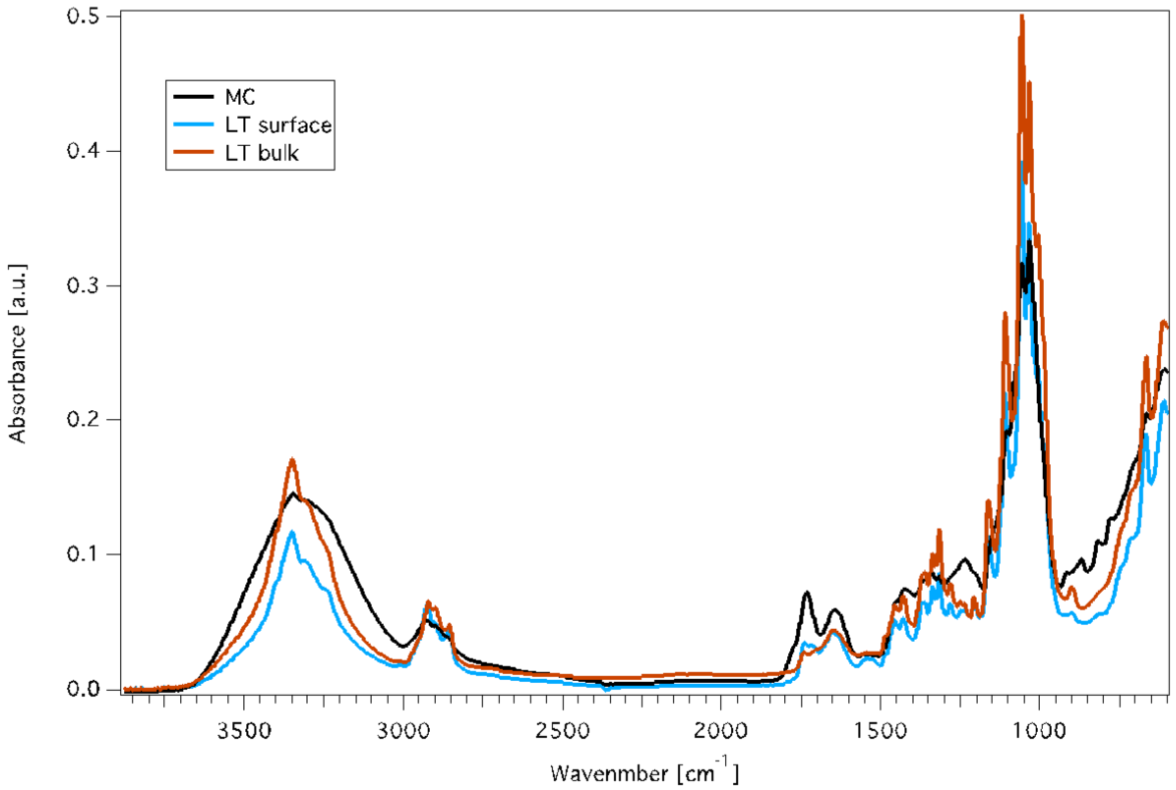


**Figure S5**

Fourier Transform Infrared Spectroscopy (FTIR) of as fabricated (MC) and lecithin tanned (LT) microbial cellulose, showing chemical changes are observed throughout the bulk of the biotextile. Bulk data was obtained by slicing hydrated LT samples in half before drying and exposing the face of the sample corresponding to the bulk region to the FTIR beam.

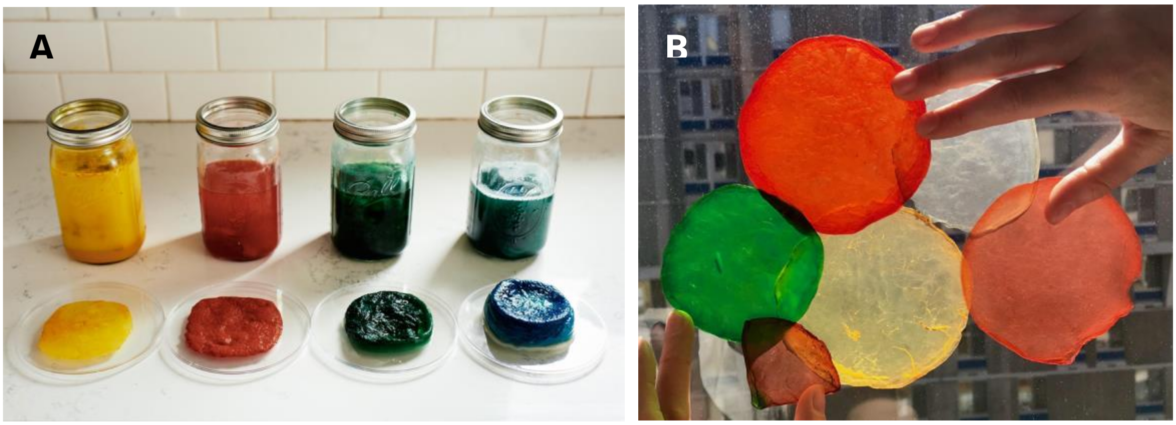


**Figure S6**

**A)** Examples of MC pellicles with color incorporated during microbial biosynthesis, using turmeric (yellow), cochineal (red), chlorophyll (green), and blue food coloring. **B)**: Examples of naturally dyed MC biotextiles after dehydration, including a transparent white obtained by immersion of MC in 0.5 M NaOH for one hour, followed by rinse in distilled water until reaching a neutral pH. See methods for details.

**A**

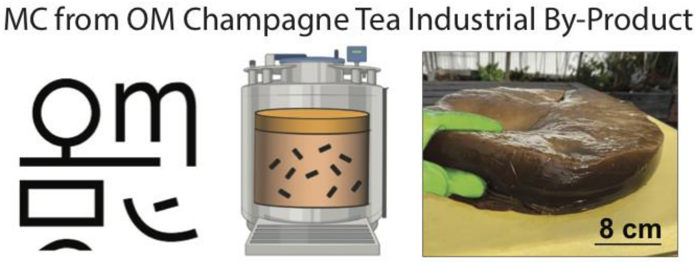


**B**

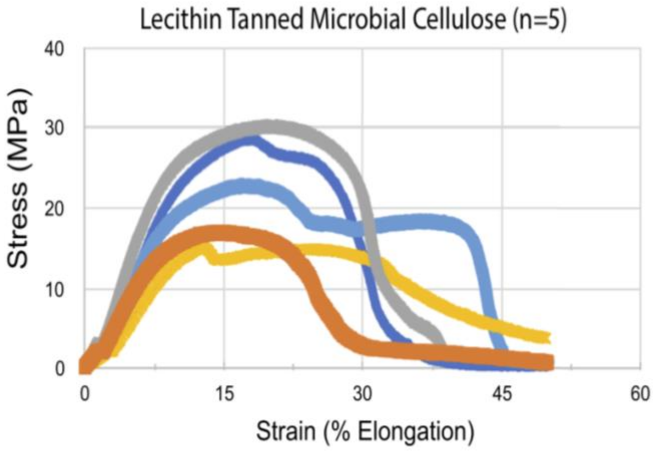


**C**

| Sample | Thickness (mm) | Width (mm) | Area ($mm^2$) | Young's Modulus (MPa) | Max Stress (MPa) | Yield Strength (MPa) | Resilience (MPa) | Toughness (MPa) |
|---|---|---|---|---|---|---|---|---|
| 1 | 1.55 | 12.70 | 19.69 | 213.71 | 17.13 | 12.21 | 0.37 | 3.75 |
| 2 | 1.38 | 12.70 | 17.53 | 341.44 | 30.21 | 20.59 | 0.67 | 7.69 |
| 3 | 1.60 | 12.70 | 20.32 | 172.59 | 15.17 | 12.34 | 0.47 | 4.95 |
| 4 | 1.52 | 12.70 | 19.30 | 271.54 | 22.95 | 15.50 | 0.49 | 7.44 |
| 5 | 1.19 | 12.70 | 15.11 | 298.06 | 28.90 | 19.44 | 0.70 | 6.39 |
| Avg | 1.45 | 12.70 | 18.39 | 259.47 | 22.87 | 16.02 | 0.54 | 6.04 |
| STD | 0.17 | 0.00 | 2.1 | 67.08 | 6.76 | 3.90 | 0.14 | 1.67 |

**Figure S7**

**A)** Microbial cellulose obtained as a by-product from commercial kombucha fermentation, provided by OM Champagne Tea, Mount Kisco, NY, corresponding to biotextile 2 in Figure 6 and the LCA analysis. Mechanical tensile data (n=5) after lecithin tanning and drying under ambient conditions expressed as: **B)** stress-strain curves; and **C)** specific parameters including modulus, max strength, yield strength, resilience and toughness.

| CS (crystallite size (nm)) L[200] | | | | | CS (crystallite size (nm)) L[101] | | | |
|---|---|---|---|---|---|---|---|---|
| n | MC | LT | LTS | Smoked | MC | LT | LTS | Smoked |
| 1 | 6.371 | 6.652 | 6.489 | 7.482 | 4.694 | 4.970 | 4.937 | 5.006 |
| 2 | 6.702 | 6.646 | 6.571 | 5.826 | 4.777 | 4.943 | 4.987 | 4.852 |
| 3 | 6.472 | 7.165 | 6.349 | 5.912 | 4.774 | 4.927 | 4.853 | 4.844 |
| 4 | 6.559 | 6.517 | 6.276 | 6.327 | 4.751 | 4.870 | 4.910 | 4.827 |
| 5 | 6.604 | 6.433 | 6.514 | 7.380 | 4.824 | 4.949 | 4.867 | 4.958 |
| **Average:** | **6.542** | **6.683** | **6.440** | **6.585** | **4.764** | **4.932** | **4.911** | **4.897** |
| STD | 0.127 | 0.285 | 0.123 | 0.796 | 0.047 | 0.038 | 0.054 | 0.080 |

| n=5 | d-spacing [200] (nm) | d-spacing [101] (nm) |
|---|---|---|
| **MC** | 0.60± 0.003 | 0.39 ± 0.001 |
| **LT** | 0.61 ± 0.001 | 0.39 ± 0.001 |
| **LTS** | 0.60 ± 0.031 | 0.39 ± 0.0011 |
| **S** | 0.60 ± 0.008 | 0.39 ± 0.0034 |

**Table S1**

Crystallite size (CS) and d-spacing in nm along the [200] and [101] planes for microbial nanocellulose before and after lecithin and aldehyde treatment, obtained from X-ray diffraction (XRD) Bragg peaks, where $d_i$ is the initial lattice spacing, $d_f$ is the final lattice spacing, with d calculated according to Bragg's Law: $\lambda = 2d_{hkl}*\sin\theta$ and $CS=k\lambda/(FWHM*\cos\theta)$, where k=0.9 is the correction factor, λ=1.54056 Å is the x-ray wavelength, FWHM is the full-width at half maximum of the Bragg peak, and θ is the diffraction angle, as described in Methods.

| Peak Intensities [a.u.] | MC | LT surface | LT bulk |
|---|---|---|---|
| $A_{897}$ | 0.066 | 0.067 | 0.077 |
| $A_{1375}$ | 0.068 | 0.072 | 0.086 |
| $A_{1430}$ | 0.056 | 0.061 | 0.072 |
| $A_{2917}$ | 0.081 | 0.068 | 0.067 |

| IR Crystallinity Ratio | MC | LT surface | LT bulk |
|---|---|---|---|
| TCI ($A_{1372}/A_{2900}$) | 0.840 | 1.059 | 1.280 |
| LOI ($A_{1429}/A_{897}$) | 0.849 | 0.911 | 0.935 |
| **Absorbance Ratio ($A_{2917}$)** | | | |
| $A_{897}$ | 0.815 | 0.985 | 1.149 |
| $A_{1375}$ | 0.840 | 1.058 | 1.284 |
| $A_{1430}$ | 0.691 | 0.897 | 1.075 |

**Table S2**

Total Crystallinity Index (TCI), Lateral Order Index (LOI) and absorbance ratio for the peaks relative to $A_{2917}$ used to probe hydrogen bond intensity (HBI) obtained from FTIR data. The data shows a slight in crystallinity, order, and decrease in hydrogen bonding due to formation of chemical cross-links (phosphates, methylene and carboxyl groups) with lecithin tanning (LT compared to as-fabricated MC) throughout the bulk of the samples.

| Impact category | Normalization | Unit | Weighting (%) |
|---|---|---|---|
| Ozone depletion | 1.61E-01 | kg $CFC^{-11}$ eq /year /capita | 2.40E+00 |
| Global warming | 2.42E+04 | kg $CO_2$ eq /year /capita | 3.49E+01 |
| Smog | 1.39E+03 | kg $O_3$ eq /year /capita | 4.80E+00 |
| Acidification | 9.09E+01 | kg $SO_2$ eq /year /capita | 3.60E+00 |
| Eutrophication | 2.16E+01 | kg N eq /year /capita | 7.20E+00 |
| Carcinogenics | 5.07E-05 | CTUh /year /capita | 9.60E+00 |
| Non-carcinogenics (Hormonally active agents (HAA)) | 1.05E-03 | CTUh /year /capita | 6.00E+00 |
| Respiratory effects | 2.43E+01 | kg PM2.5 eq /year /capita | 1.08E+01 |
| Ecotoxicity | 1.10E+04 | CTUe /year /capita | 8.40E+00 |
| Fossil fuel depletion | 1.73E+04 | MJ surplus /year /capita | 1.21E+01 |

**Table S3**
LCA normalization and weighting factors used to convert specific impacts to an impact factor expressed in mPts per functional unit (here 1 $m^2$ of textile).

| **Textile** | **Leather (By-product)** | **Leather (Co-product)** | **PU textile** | **Cotton** | **Biotextile$_{LAB}$** | **Biotextile$_{BP}$** |
|---|---|---|---|---|---|---|
| Mass/m$^2$ (kg) | 0.8 | N/A (inputs/m$^2$) | 0.66 | 0.25 | N/A (inputs/m$^2$) | N/A (inputs/m$^2$) |
| Material inventory | Cow hides, chromium process | Cow hides, slaughter house, chromium process | Table S5 | Table S6 | Table S7 | Table S8 |
| **Total Impacts (mPts)** | **3.82** | **989** | **0.62** | **0.47** | **0.30** | **9.63E-02** |
| Acidification | 2.1E-03 | 0.62 | 3.2E-02 | 2.32E-02 | 2.74E-03 | 5.75E-04 |
| Ecotoxicity | 1.5E-02 | 39.1 | 3.1E-02 | 0.06 | 4.88E-02 | 2.05E-02 |
| Eutrophication | 3.9E-04 | 3.36 | 3.5E-02 | 3.66E-02 | 5.55E-02 | 5.06E-02 |
| Global warming | **1.7E-02** | **1.85** | **0.17** | **9.37E-02** | **1.3E-02** | **5.29E-03** |
| Ozone depletion | 1.9E-05 | 8.0E-04 | 1.3E-04 | 2.38E-05 | 1.75E-05 | 8.28E-06 |
| Fossil fuel depletion | 1.4E-02 | 0.64 | 0.12 | 1.60E-02 | 9.42E-03 | 4.09E-03 |
| Carcinogenics | **3.63** | **937** | **0.16** | **0.146** | **6.85E-02** | **7.07E-03** |
| Non-carcinogenics (HAA) | 2.8E-03 | 5.78 | 2.4 E-02 | 5.11E-02 | 8.75E-02 | 7.00E-03 |
| Respiratory effects | 1.4E-03 | 0.6 | 1.0 E-02 | 2.52E-02 | 6.67E-03 | 1.00E-03 |
| Smog | 1.7E-03 | 0.19 | 4.5E-02 | 1.67E-02 | 2.80E-03 | 8.06E-05 |
| **$CO_2$ eq. (kg)** | **1.17** | **128** | **11.50** | **6.50** | **0.89** | **0.37** |

**Table S4**
**Cradle to Gate LCA - Manufacturing Impacts for 1 m$^2$ of textile.** Values obtained from LCA based within the Sustainable Minds software using the Ecoinvent data sets described.

| Input (mass) | Polyester (PET) fabric (594 g) | Epoxy resin (54.9 g) | Ethyl acetate (9.9 g) | Citric acid (1 g) | DMF (11.55 g) |
|---|---|---|---|---|---|

| | | | | | |
|---|---|---|---|---|---|
| **Total impacts** | **0.58** | **3.4E-02** | **2.39E-03** | **6.0E-04** | **9.81E-03** |
| Acidification | 3.08E-02 | 1.04E-03 | 5.29E-05 | 1.70E-05 | 5.86E-05 |
| Ecotoxicity | 2.86E-02 | 2.34E-03 | 1.02E-04 | 1.18E-04 | 1.04E-04 |
| Eutrophication | 2.60E-02 | 8.86E-04 | 1.32E-04 | 3.38E-05 | 7.8E-03 |
| Global warming | 0.159 | 5.76E-03 | 4.62E-04 | 8.76E-05 | 3.87E-04 |
| Ozone depletion | 1.27E-04 | 1.52E-08 | 6.10E-07 | 1.16E-07 | 6.76E-07 |
| Fossil fuel depletion | 0.113 | 6.33E-03 | 6.53E-04 | 3.32E-05 | 4.89E-04 |
| Carcinogenics | 0.145 | 1.1E-02 | 7.42E-04 | 2.15E-04 | 7.45E-04 |
| Non-carcinogenics /HAA | 2.10E-02 | 3.02E-03 | 1.36E-04 | 5.64E-05 | 1.32E-04 |
| Respiratory effects | 8.41E-03 | 1.76E-03 | 4.62E-05 | 2.46E-05 | 4.85E-05 |
| Smog | 4.30E-02 | 1.78E-03 | 4.94E-05 | 1.43E-05 | 4.25E-05 |
| **$CO_2$ eq. (kgs)** | **11.0** | **0.4** | **0.032** | **6.1E-03** | **2.68E-02** |

**Table S5**
Impacts for 1 m$^2$ of polyurethane (PU)-coated polyester textile (synthetic leather) by input computed within the LCA methodology (Figure 6).

| Input | Fiber production | Yarn production | Weaving process |
|---|---|---|---|
| **Total impacts** | **0.13** | **0.15** | **0.19** |
| Acidification | 2.56E-03 | 8.96E-03 | 1.16E-02 |
| Ecotoxicity | 4.47E-02 | 6.23E-03 | 7.62E-03 |
| Eutrophication | 3.35E-02 | 1.34E-03 | 1.73E-03 |
| Global warming | 9.0E-03 | 3.76E-02 | 4.71E-02 |
| Ozone depletion | 8.22E-06 | 7.55E-06 | 8.01E-06 |
| Fossil fuel depletion | 4.47E-03 | 5.44E-03 | 6.12E-03 |
| Carcinogenics | 1.65E-02 | 5.85E-02 | 7.1E-02 |
| Non carcinogenics | 1.67E-02 | 1.5E-02 | 1.94E-02 |

| | | | |
|---|---|---|---|
| Respiratory effects | 2.13E-03 | 9.93E-03 | 1.32E-02 |
| Smog | 2.0E-03 | 6.35E-03 | 6.35E-03 |
| **$CO_2$ eq. (kgs)** | **0.625** | **2.61** | **3.27** |

**Table S6**

Impacts for 1m$^2$ of cotton fabric (250 g/m$^2$) by input computed within the LCA methodology in Sustainable Minds Software.

| **Input** | **Sugar** | **Sunflower seed oil** | **Green tea** | **Natural gas** | **Tap water** |
|---|---|---|---|---|---|
| **Inventory** | 1.74 kg | 296 ml | 120 g | 1744 Btu | 40 l |
| **Total Impacts** | **0.192** | **1.92E-02** | **1.24E-04** | **1.04E-02** | **1.34E-03** |
| Acidification | 2.08E-03 | 5.17E-04 | 1.42E-06 | 1.04E-04 | 4.32E-05 |
| Ecotoxicity | 2.82E-02 | 2.05E-02 | 5.50E-05 | 4.99E-05 | 6.85E-05 |
| Eutrophication | 4.85E-03 | 5.06E-02 | 1.15E-05 | 3.4E-05 | 8.51E-06 |
| Global warming | 5.08E-03 | 3.31E-03 | 2.95E-06 | 4.23E-03 | 2.51E-04 |
| Ozone depletion | 5.12E-6 | 4.92E-6 | 2.43E-09 | 7.38E-06 | 1.29E-07 |
| Fossil fuel depletion | 2.65E-03 | 1.91E-03 | 1.40E-06 | 4.79E-03 | 7.28E-05 |
| Carcinogenics | 6.04E-02 | 6.49E-03 | 1.95E-05 | 8.77E-04 | 7.17E-04 |
| Non-carcinogenics /HAA | 8.04E-02 | 6.93E-03 | 3.08E-05 | 1.01E-04 | 1.13E-04 |
| Respiratory effects | 5.61E-03 | 9.64E-04 | 1.21E-06 | 6.07E-5 | 3.65E-05 |
| Smog | 2.61E-03 | 1.11E-06 | 5.55E-07 | 1.59E-04 | 3.10E-05 |
| **$CO_2$ eq. (kg)** | **0.353** | **0.229** | **2.05E-02** | **0.294** | **1.74E-02** |

**Table S7**

Cradle to Gate LCA -Biotextile$_{LAB}$- Impacts for laboratory scale biosynthesis and lecithin tanning of 1 $m^2$ of LT MC textile by input.

| Input | Tap water | Natural gas | Sunflower seed oil |
|---|---|---|---|
| **Inventory** | 10 l | 3864.68 Btu | 296 ml |
| **Total Impacts** | **3.35E-04** | **4.71E-03** | **9.12E-02** |
| Acidification | 1.08E-05 | 4.67E-05 | 5.17E-04 |
| Ecotoxicity | 1.71E-05 | 2.25E-05 | 2.05E-02 |
| Eutrophication | 2.13E-06 | 1.54E-05 | 5.06E-02 |
| Global warming | 6.27E-05 | 0.00192 | 3.31E-03 |
| Ozone depletion | 3.22E-08 | 3.33E-06 | 4.92E-6 |
| Fossil fuel depletion | 1.82E-05 | 2.16E-03 | 1.91E-03 |
| Carcinogenics | 1.79E-04 | 3.96E-04 | 6.49E-03 |
| Non-carcinogenics /HAA | 2.82E-05 | 4.57E-05 | 6.93E-03 |
| Respiratory effects | 9.12E-06 | 2.74E-05 | 9.64E-04 |
| Smog | 7.75E-06 | 7.17E-05 | 1.11E-06 |
| **$CO_2$ eq. (kg)** | **4.35E-03** | **0.133** | **0.229** |

**Table S8**

Cradle to Gate LCA -Biotextile$_{BP}$- Impacts for 1 m$^2$ of biotextile fabricated using microbial nanocellulose obtained as SCOBY by-product of commercial food and (kombucha) beverage production, by input.

| Sample (n=5) | Initial mass (g) Day 0 | Final mass (g) Day 60 | Mass loss (%) | Average mass loss (%) |
|---|---|---|---|---|
| MC 1 | 0.09 | 0.024 | 73.33 | |
| MC 2 | 0.09 | 0.02 | 77.78 | |
| MC 3 | 0.09 | 0.025 | 72.22 | |
| MC 4 | 0.09 | -- | -- | |
| MC 5 | 0.09 | -- | -- | |
| **MC** | | | | **74.45 +/- 2.94** |
| LT 1 | 0.05 | 0.013 | 74 | |
| LT 2 | 0.04 | 0.012 | 70 | |
| LT 3 | 0.04 | 0.009 | 77.5 | |
| LT 4 | 0.04 | 0.008 | 80 | |
| LT 5 | 0.04 | 0.008 | 80 | |
| **LT** | | | | **76.30+/- 4.30** |

**Table S9**
Biodegradation in natural terrestrial environment of 2 x 2 cm samples (n=5) of as fabricated (MC) and lecithin tanned (LT) microbial biotextiles evaluated as mass lost after 60 days outdoors in soil. Two of the MC samples were not found upon retrieval. The slightly larger mass loss for LT is likely due to the lower density (thickness of the MC nanofiber mesh) of the samples relative to MC (0.0105 $g/cm^2$ vs 0.0225 $g/cm^2$).

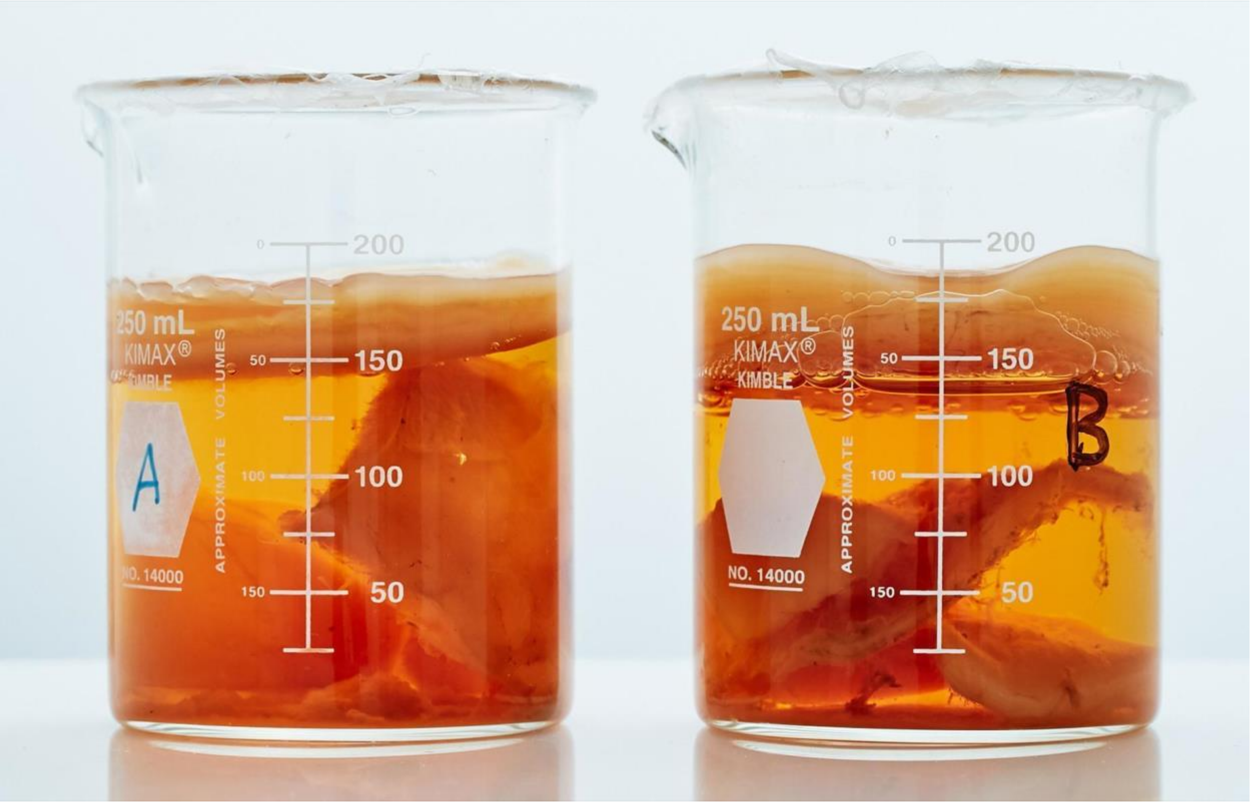


Microbial Cellulose Biosynthesis Time Lapse

**Movie S1**

Time lapse of microbial cellulose biosynthesis over 21 days at 24°C using: **A)** sucrose and **B)** fructose as carbon sources shows tunable and scalable biosynthesis highlighting the ability of *G. Xylinum* to utilize a variety of nutrient sources in its carbon metabolism.

**Movie S2**

Water repellency of waxed MC biotextile
